# Sparse mesenchymal cell networks as a fluid under tension

**DOI:** 10.64898/2025.12.07.692626

**Authors:** Alex T. Grigas, Gabriella Shih Ping Hsia, Rajendra Singh Negi, Eirini Maniou, Gabriel L. Galea, Arthur Michaut, Alessandro Mongera, M. Lisa Manning

**Affiliations:** Department of Physics and BioInspired Institute, Syracuse University, Syracuse, New York 13244, USA; Department of Cell and Developmental Biology, University College London, London WC1E 7JE, UK; School of Biotechnology, Dublin City University, Dublin D09 K20V, Ireland; Department of Developmental Biology and Cancer, University College London Great Ormond Street Institute of Child Health, London WC1N 1 EH, UK; Department of Developmental and Stem Cell Biology, Institut Pasteur, Université de Paris Cité, CNRS UMR 3738, 75015 Paris, France

## Abstract

Sparse mesenchymal cellular networks are ubiquitous across animals, shaping both embryonic and adult structures through dynamic interactions with epithelia. Yet, the physical principles underlying their collective behaviors remain elusive, as their stellate cells and large extracellular spaces—filled with matrix or interstitial fluid—pose significant experimental and computational challenges. Here, we demonstrate that the avian presomitic mesoderm (PSM), a canonical embryonic mesenchymal tissue, behaves as a fluid under tension, exhibiting structural organization that cannot be explained by simple Brownian-like cell motion. Through quantitative modeling, we identify contact inhibition of locomotion (CIL)—where cells actively retract and move away upon contact—as a key mechanism that enables sparse mesenchymal networks to sustain macroscopic tension while flowing like a fluid. Consistent with this mechanism, colliding PSM cells turn away from each other, and spreading PSM explants accelerate outward as interior voids open. Simple continuum equations relate observable cell-scale parameters to the emergent remodeling dynamics observed in both experiments and simulations. Together, these findings put forward an unrecognized mechanical role for CIL, extending its influence beyond collective migration, and establish the fluid-under-tension state as a distinct class of tissue behavior that describes key developing embryonic tissues and may illuminate how matrix-rich adult tissues fluidize during tumorigenesis.

---

During embryonic morphogenesis – and its reactivation in regeneration and cancer progression – biological tissues must alternate between flowing like fluids to change shape and resisting flow like solids to maintain structural integrity [1]. The mechanisms that enable these fluid–solid transitions are now well characterized in certain tissue types, such as confluent epithelial layers [2–6] or tissues with bubble-shaped cells that generate foam-like architectures [7, 8]. In contrast, much less is known about stellate mesenchymal tissues, whose cells exhibit highly irregular shapes with protrusions and large extracellular gaps [9–11].

Here, we use the presomitic mesoderm (PSM)—the engine driving posterior body axis elongation [7, 9, 12]—as a model to investigate the mechanical regulation of fluidity in sparse mesenchymal tissues composed of star-like cells. We focus on the avian PSM, which forms a highly porous cellular network with irregular gaps filled by soft extracellular matrix (ECM) components such as hyaluronic acid (HA) [9].

The highly porous nature of the cell network in mesenchymal tissues is important for its biological function. Recent work on the avian PSM has shown that the large intercellular spaces are essential for body-axis elongation, as elongation is driven by the expansion of the extra-cellular volume [9]. Defects in mesenchymal tissue mor-phogenesis have been implicated in congenital disorders such as Robinow syndrome, in which impaired cell rear-rangements result in reduced length of several embryonic structures, including the limbs, mandibular arches, and vertebral column [13–18]. Molecularly, a major regulator of PSM fluidity is Fibroblast Growth Factor (FGF) signaling, which activates cell motility through ERK [12, 19] and modulates extracellular expansion through HA synthesis [9].

Previous work has highlighted two distinct mechanical pathways that govern tissue fluidity in other tissue types. In confluent tissues such as epithelial monolayers—where polygonal cells form a continuous sheet without gaps or overlaps—the preferred cell shape, determined by mechanical parameters such as cortical tension and cell–cell adhesion, is the principal factor controlling tissue fluidity [2–6]. Vertex models [20] have successfully reproduced these behaviors, revealing that fluid-to-solid transitions correspond to a second-order rigidity transition driven by changes in geometric properties rather than the network topology. [21–24].

In non-confluent tissues with bubble-shaped cells, such as the zebrafish PSM, fluid–solid transitions are instead controlled by the topology of cell–cell contacts [7]. In these systems, increasing the extracellular volume fraction or reducing cell–cell adhesion lowers the packing fraction and decreases the average number of contacts per cell. Soft-sphere and bubble models [25, 26] have successfully captured these behaviors [7, 8], showing that the transitions between fluid and solid states arises from a first-order rigidity, or jamming, transition.

Notably, despite their homology, the zebrafish PSM displays distinct architectural features compared with its avian counterpart: cells are more densely packed, and the ECM has been proposed to play only a minor role [27]. As a consequence, the zebrafish PSM is well-described by soft-sphere models [7]. Sphere-based models also capture certain large-scale behaviors of the avian PSM, such as the emergence of tissue expansion driven by gradients in cell motility [9]. Yet, such models are unlikely to mechanistically predict other emergent features of tissue mechanics, as their assumptions about how contacts scale with packing fraction fail for the highly irregular geometries found in stellate mesenchymes.

Similarly, vertex models do not capture the porous nature of the stellate cellular network, and yet they also seem to work well to predict features at the tissue scale. For example, mesenchymal tissues have been shown to generate localized structures such as dermis follicles or intestinal villi through a dewetting transition driven by tissue scale tension [28, 29]. These behaviors have been successfully captured using confluent vertex models [29], or continuum contractile fluid models [28]. In addition, work on the avian PSM has demonstrated that it also behaves as a fluid on long timescales [12], and confluent vertex and cellular Potts models can also capture features of its elongation [30, 31].

Thus, an important open question is how the porous, highly non-confluent cellular structure of stellate mesenchymal tissues gives rise to the emergent, tissue-scale features that seem to require both tension and fluidity at the same time. Is there direct evidence for these tissues being under tension despite their fluid-like nature? If so, how is that possible, given that tension should presumably lead to density-increasing instabilities in the porous cellular network, and do the cells themselves show the behavior needed to prevent them?

## Avian PSM tissue is fluid-like, under tension and uniformly textured

First, we demonstrate the unique architectural properties of the avian PSM. Fig. 1(a), highlights the position of the PSM relative to the postcranial body axis. Zooming in, Fig. 1(b) demonstrates the porous nature of the PSM cellular network, especially compared to the neighboring notocord (NC) and intermediate/lateral plate mesoderm (IM/LPM). At the cellular scale, we further show the stellate shapes of the PSM in Fig. 1(c). By injecting labeled dextran into the intercellular space of a membrane-labeled PSM, we can then visualize both the cellular network and the porous interstitial space. (See Fig. 1(d)). To better represent the shape of individual cells, we show in Fig. 1(e) a time course of a cell within a mosaically labeled PSM, which emphasizes the dynamic nature of the protrusions extended from these stellate cells.

**FIG 1.**
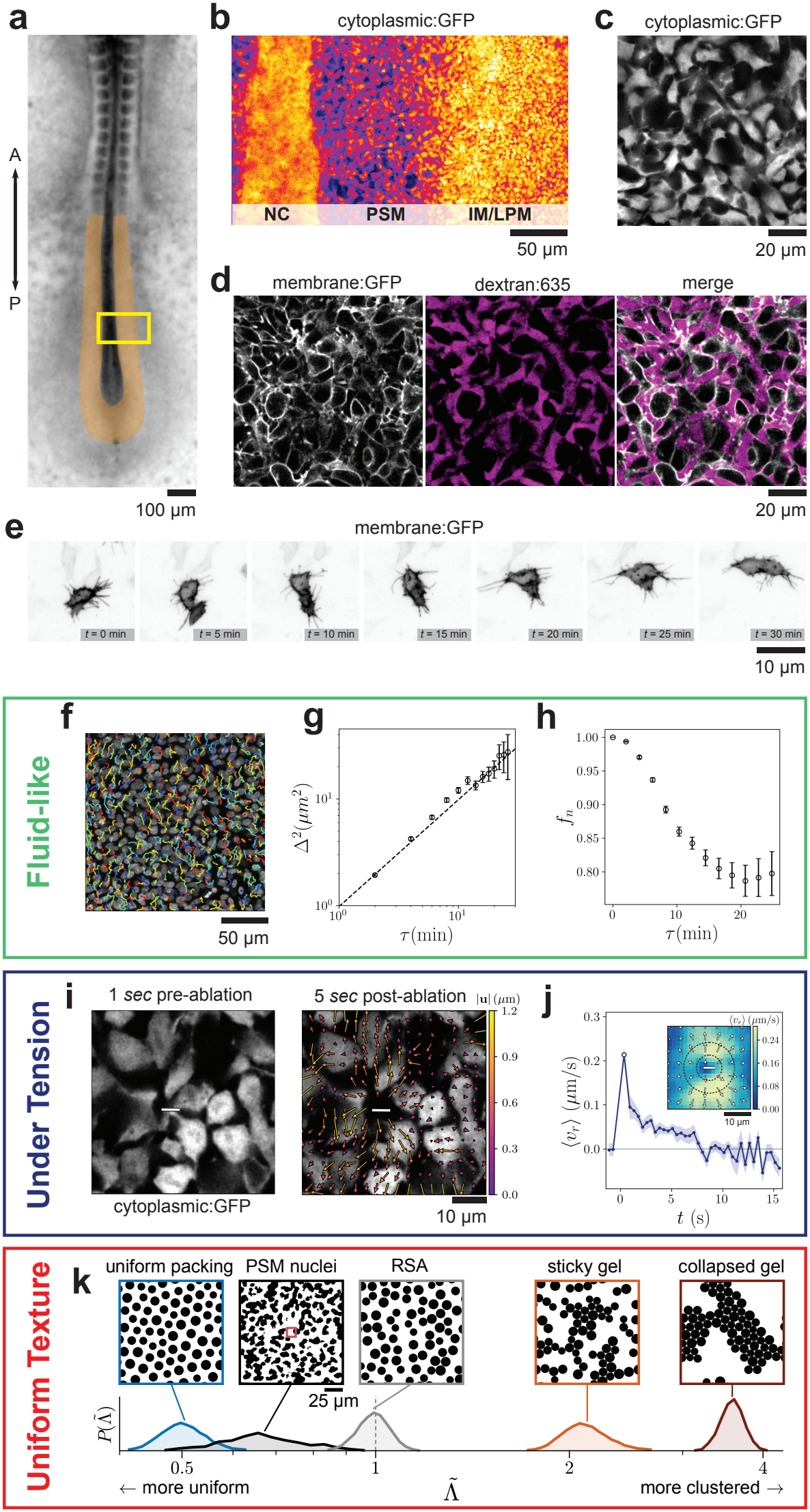
The avian PSM is a uniform fluid under tension. (a) Brightfield image of the post-cranial body axis of an avian embryo at stage HH XII. The PSM is pseudo-colored in orange. (b) Zoomed in view of the rectangular inset annotated in (a) showing the medio-lateral organization of notocord (NC), presomitic mesoderm (PSM) and intermediate/lateral plate mesoderm (IM/LPM). (c) Network of PSM cells labeled with a cytosplamic GFP. (d) Membrane-labeled cellular network with dextran injected into the extracellular space. (e) Mosaically membrane-labeled PSM cell over 30 min. (f) Nuclei tracking over a 2 hour window with time axis color-coded from blue to red. (g) Mean squared distance difference between neighbors Δ^2^ vs lag time *τ*. The dashed line indicates the diffusive slope of 1. (h) Fraction of neighbors that remain within the neighbor cutoff distance *f*_*n*_ vs lag time *τ*. Error bars in (g) and (h) indicate the standard error. (i) Cytoplasmic:GFP cell network before and after laser ablation. The grey rectangle indicates the ablation site. (j) The average retraction velocity from PIV analysis vs time post-ablation, averaged over *n* = 25 ablations across 7 embryos. Error bars indicate standard error. Inset: Average PIV retraction field at the peak retraction velocity. Radial retraction magnitude is color-coded from blue to yellow. (k) Probability distributions and example images of the range of cell-scale textures 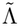 on a log-scale. The PSM nuclei (black inset) lie between highly uniform diluted jammed packings (blue) and RSA (grey). Quenching attractive particles with a short range (orange) and long-range (brown) interactions result in increasingly clustered gels.

Next, we demonstrate that during elongation the avian PSM behaves as a fluid under tension *in vivo*, with the cells simultaneously exhibiting three distinct properties:

1) cells diffuse and exchange neighbors like a fluid, 2) the cellular network of the PSM is under tension, and 3) cells in the PSM are uniformly dispersed with substantial gaps.

First, we track nuclei in the PSM over 2 hours using Ultrack [32]. Nuclear labeled cells and a portion of cell tracks are shown in Fig. 1(f). As the PSM is undergoing elongation, cell tracks are both moving along the body axis (drift) and moving relative to each other (diffusion), which both contribute to the mean squared displacement. To remove drift and quantify solely diffusion, we measure instead the mean-squared distance difference [33] between cells that were initially neighbors,

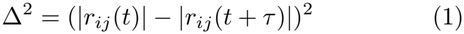

where |*r*_*ij*_(*t*)| is the distance between initially neighboring cells *i* and *j* at time *t* and lag-time *t* + *τ*, averaging over all initial neighbors and all time origins *t*. Neighboring cells are defined as nuclei that are initially within |*r*_*ij*_(*t*)| ≤ 12 *µ*m. In Fig. 1(g), we show that Δ^2^ grows linearly, indicating the motion without the elongation drift is diffusive. This diffusive motion agrees with previous measurements of cell motility in the avian PSM [12, 34], which was interpreted as a “random cell walk”. However, diffusive statistics do not require random motion, as we show below. We also quantify the fraction of initial neighbors |*r*_*ij*_(*t*)| ≤ 12 *µ*m that have moved away beyond a threshold distance |*r*_*ij*_(*t* + *τ*)| *>* 15 *µ*m after a lag-time *f*_*n*_(*τ*). After 10 minutes, a substantial fraction (~15%) of neighbors have exchanged, shown in Fig. 1(h). We find similar results for other threshold distances (see Supplemental Information, SI).

Second, to demonstrate cell-cell tension, we perform laser ablation to ablate single cell-cell contacts as shown in Fig. 1 (i) (*n* = 25 cuts across 7 embryos). Z-stacks acquired immediately after each cut confirm that ablations target the PSM, approximately 20–30 µm from the ventral surface, while the overlying endoderm remains intact (see SI). Using PIV analysis, we quantify radial retraction velocities of the tissue away from the ablation site [35]. (See Fig. 1 (j)). There is a clear and significant retraction velocity, which peaks within 1 second at ~0.2 *µm/s* and decays within 10 seconds, similar to the retraction velocity reported after severing single tensioned cell–cell junctions in the *Drosophila* germband [36–38]. Additionally, a dipole-like structure can be seen in the average retraction velocity field (Fig. 1 (j) inset), consistent with the expected response in cutting a connection in a tensioned elastic network.

Lastly, we develop a method to quantify the spatial dispersion of cells in the PSM, termed the ‘cell-scale texture’, as we observe that cells remain remarkably uniformly dispersed despite high tensions and rearrangement dynamics. First, we binarize the nuclear labeled PSM microscopy images, shown in Fig. 1 (k). We measure the average pixel intensity *λ*, i.e., the fraction of the window covered by nuclei, inside a small square window of side *r* approximately the size of a cell. At this level, for a highly clumped sample, there will be many windows that are entirely full when passed over the clumps, and many windows that are empty as they pass over void regions. Because of this, there is a high variance in the average pixel intensity as the window is passed over the material. In contrast, in a very uniform material, one will find that the average pixel intensity in the local window is approximately the same across the entire image and therefore the variance is small. Therefore, we term the variance of the windowed pixel intensity the cell-scale texture Λ. (In materials science, the same quantity is known as the local volume-fraction variance [39]. The scaling of Λ vs *r* in much larger windows also quantifies hyperuniformity, which we do not consider here as the PSM is only ~ 100 *µ*m across [40, 41]).

To normalize the cell-scale texture Λ, we use the same measure, the particle-scale texture, for randomly placed non-overlapping disks – random sequential addition (RSA) [42] – at the same fill fraction: 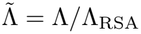. 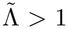 is a texture more clustered than random, such as in a sticky gel material, while 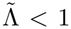 is more uniformly dispersed than random, such as in a uniform packing. See Fig. 1 (k) for example images of each type of texture. We find that the PSM nuclei data is very uniform with an average cell-scale texture 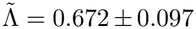. See Methods and SI for further details.

Analyzing cytoplasmically labeled PSM tissue gives a similar picture as nuclear labeling. See SI for further details.

### A minimal mesenchymal cell model

We next develop a new inter-cellular interaction model to explore whether different types of activity can result in properties observed in the PSM. To effectively model mesenchymal tissues, we need a non-confluent representation that can vary density, support tensions and undergo contact changes. We present a schematic of the hypothesized mesenchymal cell interaction in Fig. 2 (a) as four distinct interactions: i) cells far apart have no interaction, ii) if cells move closer together than a characteristic ‘repulsive contact distance’, they elastically deform and support compressive forces, iii) after cells come into contact, they form an adhesion, and as the cells are stretched apart this contact and associated cytoskeleton can support tensile forces, and iv) cells can break these adhesive contacts. In the bubble model [25], a repulsive linear spring has long been used to model the deformation in (ii); however, there is no comparably simple choice for the tensile forces in (iii), as we discuss next.

**FIG 2.**
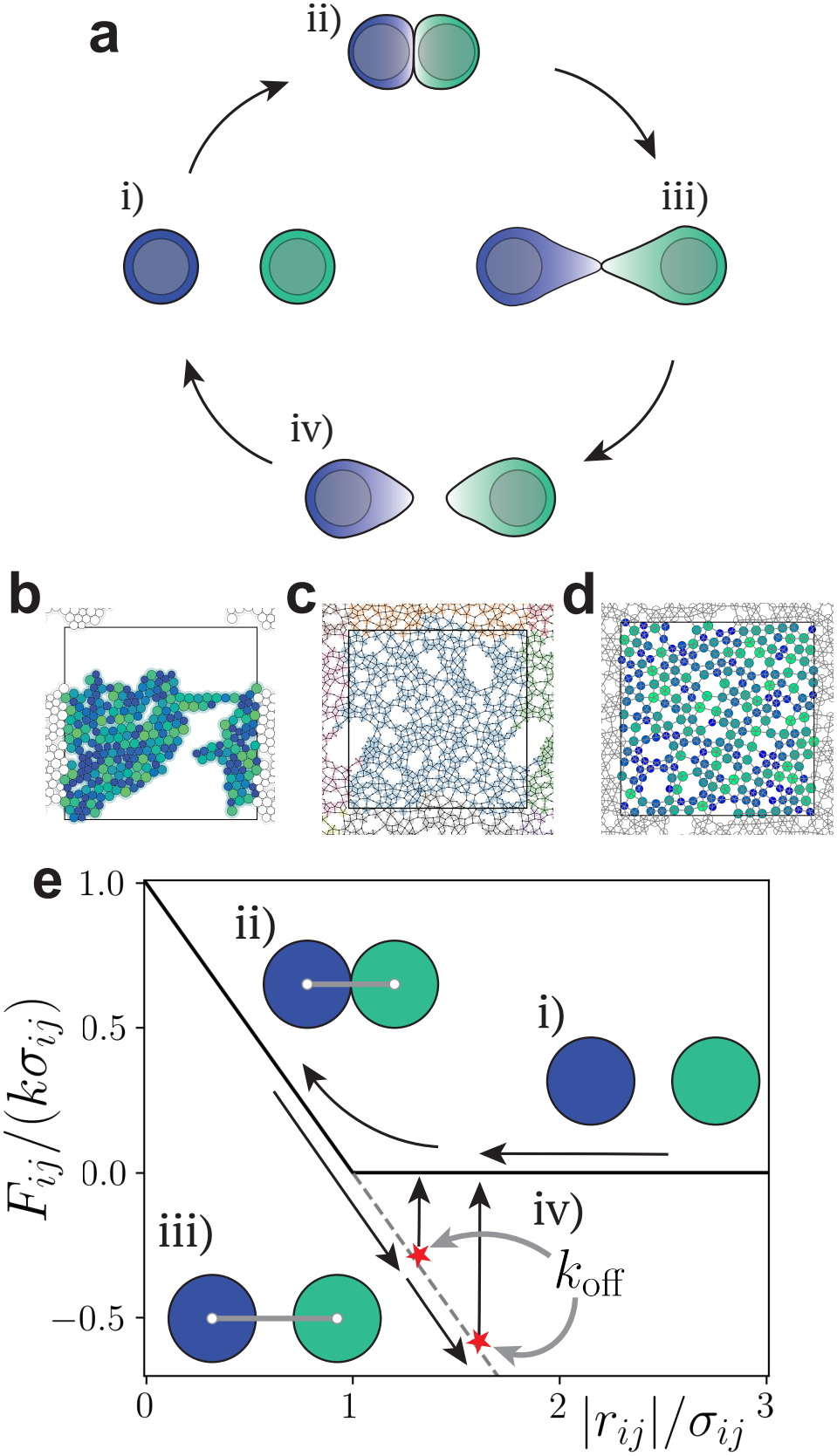
A mesenchymal cell potential. (a) A cartoon of mesenchymal cell interaction. i) At a distance, cells do not interact. ii) Cells come into repulsive contact, forming an adhesion and elastically deforming, thus generating compressive forces. iii) After forming an adhesion, cells stretched apart remain connected and deform elastically under tensile forces. iv) Cells break the adhesion and separate. (b) A packing of sticky disks under extension. Blue circles around disks indicate the attractive distance. (c) A packing of deformable patchy particles under extension. (d) A packing of hysteretic sticky particles. Colors from blue to green indicate cell size from small to large in (b) and (d). (e) Force *F*_*ij*_ vs cell-cell distance *r*_*ij*_ for the hysteretic sticky potential with the same four types of interactions as depicted in (a), where adhesions break stochastically at a constant rate *k*_off_.

To add tensile forces to the bubble model, one could simply create a sticky particle by adding an attractive contribution, which has both a repulsive distance and an attractive distance [43–45]. However, when sticky disks are compressed to form an initial network, upon extension they do not dilate but rip into gel-like clusters [46, 47]. (See Fig. 2 (b)). A detailed treatment could be achieved by an explicitly deformable particle model with attraction where the attractive energy is stronger than the cell shape deformation [48–50], although such models may not also support the strong compressive stresses arising from an internal cytoskeleton that we desire for ii).

Therefore, we next study a patchy particle, as shown in Fig. 2 (c). In this iteration, a core repulsive bubble has linear-spring arms with sticky particle adhesions at the ends [51], so that when the arm stretching energy is weaker than the adhesion strength, the network dilates uniformly upon extension, reminiscent of the avian PSM. While this iteration of the model does exhibit many features of the PSM, as we will describe in future work, it has a large number of model parameters and dynamics associated with the complex motion of the arms.

To avoid this complexity, we instead capture the key features of the PSM using a simpler repulsive bubble model with hysteretic sticking. Initially, cells behave as repulsive bubbles. Once two cells come into contact, they form an adhesion, which under extension supports tension as a double-sided linear spring. (See Fig. 2 (e)). This cell interaction results in a tensioned, uniform, dilated tissue under extension shown in Fig. 2 (d). In general, adhesive interactions in mesenchymal tissues are complex [52]. As shown in Fig 1 (d), there is significant extracellular space with soft ECM, such as HA, that can mediate cell-ECM-cell adhesive interactions, as well as direct cell-cell adhesion via molecules like N-cadherin. For simplicity, we incorporate both cell-ECM-cell and cell-cell adhesions in a single adhesive term as shown in Fig 2(a,e)(iii); future work should explore this further.

To allow fluid-like flows, the model also needs to capture remodeling of adhesive cell-cell contacts. To do so, we introduce a stochastic unbinding rate *k*_off_ which randomly removes cell-cell contacts, returning them to their initial unbound state. This uniform rate – independent of applied tension – was chosen for simplicity, and tensiondependent rates such as those found in slip or catch bonds could be added later.

The system is evolved in time using over-damped Langevin dynamics with temperature *T* = 0 to capture the viscosity of the extracellular space and isolate the cellular activity as the driving mechanism. The initial configurations are generated by quasi-static compression to a dense packing, followed by extension to the target packing fraction of *ϕ* = 0.4, a similar density to the avian PSM, although we have checked that steady state results are independent of initialization details. Unless otherwise noted, all simulations are carried out on *N* = 256 cells under periodic boundary conditions ensemble averaged over 10 independent simulations. See Methods for further details.

### CIL-like behavior is necessary to create a fluid under tension

Tension will relax as adhesive cell-cell contacts in the system are released according to *k*_off_, so a mechanism is required to generate additional tension and maintain homeostasis. In the embryo, external stretching forces involved in body-axis elongation could, in principle, supply tension globally. However, as we demonstrate in the SI, global stretching leads to clustering inconsistent with experimental observations. Instead, we consider energy to generate tension injected locally by the cells themselves. We first consider the simplest model for cellular activity, the self-propelled particle (SPP) [53]. For active Brownian particles, the active force added to the system is

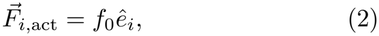

where 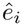 is a unit vector that diffuses randomly and cells move along 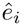 with a strength *f*_0_, resulting in a swim speed *v*_0_ = *f*_0_*/γ* where *γ* is the damping parameter [53, 54]. (See Fig. 3 (a) (left) for a schematic). We simulate the system for different combinations of *v*_0_ and *k*_off_ for a slow reorientation rate.

**FIG 3.**
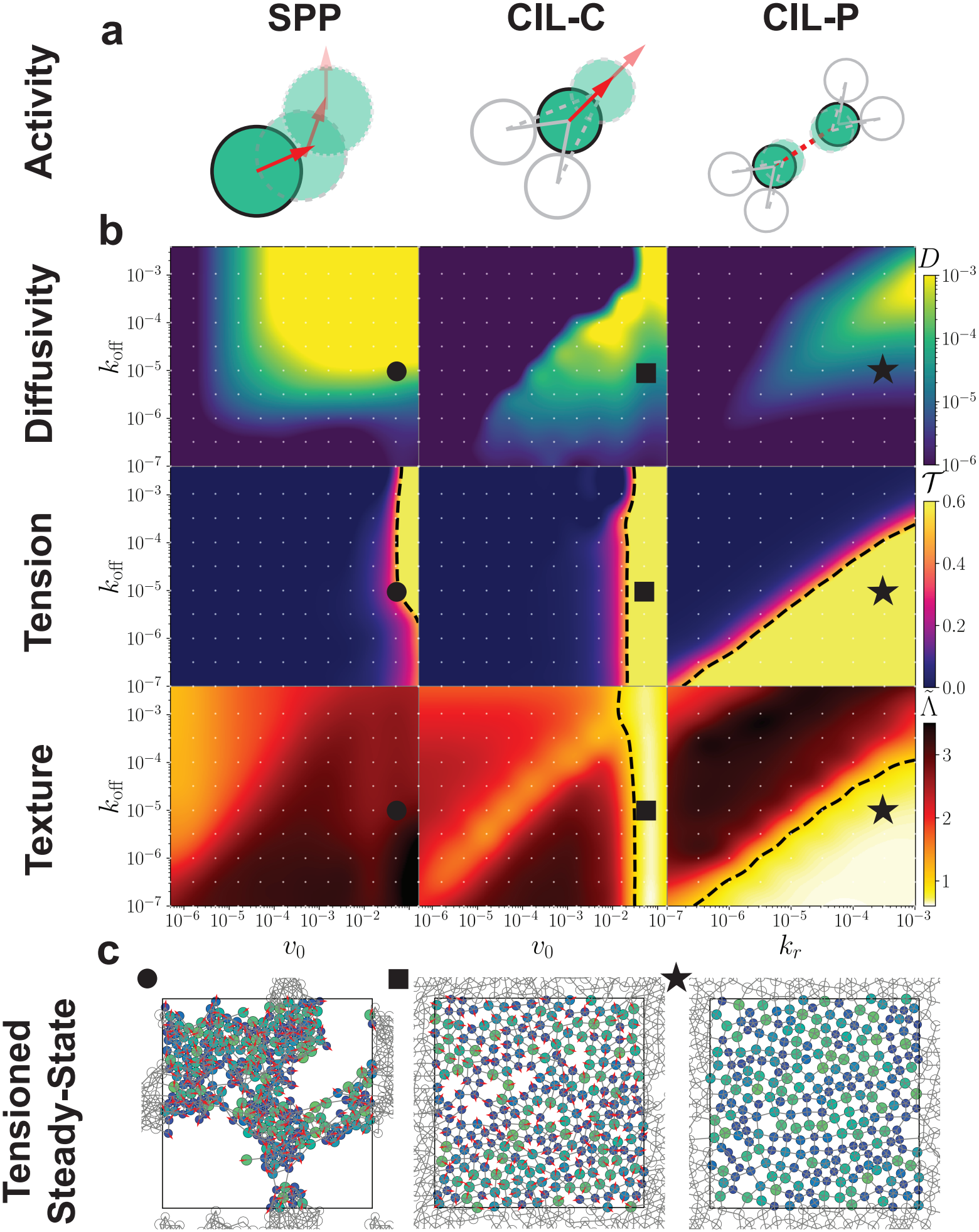
CIL is necessary to produce a tensioned fluid. (a) Schematics of the three different models for activity acting on a central particle in green. Decreased opacity indicates motion in time according to the activity. Red arrows indicate crawling activity and the red dashed line indicates the contracting cell-cell contact. (Left column) The self-propelled particle (SPP). (Center column) CIL-C active crawling model. (Right column) CIL-P active pulling model. (b) Phase diagrams. The system was evolved to reach a steady-state for a range of activities (either *v*_0_ or *k*_*r*_) and unbinding rates *k*_off_ (white points). The diffusion coefficient *D*, tension *T* and normalized cell-scale texture 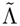 were subsequently measured with a cubic interpolation between data points. Color brightness increases from dark to yellow, and the tensioned fluid phase emerges where yellow regions overlap. Dashed lines indicate where *T >* 0.5, half the saturated tension per adhesion in CIL-P, and 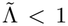. (c) Example visualizations of the final steady-state marked with a circle, square and star for SPP, CIL-C and CIL-P respectively in (b). Blue to green indicates cell size from small to large, grey lines are springs, and red arrows indicate crawling direction in SPP and CIL-C models.

We quantify the overall fluidity of the system using the diffusion coefficient *D* extracted from the mean squared displacement. Using the virial stress tensor [55], we define the tension *T* as the negative virial pressure per cellcell adhesion, the average force times separation carried by an adhesion (see Methods). We measure normalized cell-scale texture 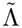 in the same manner as for the experimental images. (See Methods). In Fig. 3, we plot phase diagrams of *D, T* and 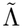. While increasing activity can generate flow and tension, the structure of the system is highly clustered in the tensioned region (Fig. 3(C)), as cells pull on one another anisotropically and gaps between cells collapse. There is no region of phase space where the random motion of the SPP model is able to generate a uniform fluid under tension.

Next, we modify how energy is injected into the system. The mode of failure in the SPP case is via clustering, caused by anisotropic tension instabilities that cause porous structures to collapse. While an interpenetrating soft ECM can exert restoring forces that counteract collapse [56], our observation that cells regularly change neighbors and reconfigure the soft matrix suggests that the ECM by itself cannot resist clustering on longer timescales.

Therefore, to generate a uniform tissue we speculate that cells must preferentially make new contacts in directions that counteract anisotropy, i.e. preferentially moving or protruding towards regions of empty space where there are no existing contacts. In cellular biology, this same behavior has long been identified in isolated cells and is referred to as contact inhibition of locomotion (CIL) [57]. We consider two different models of CIL. First, we consider active crawling (CIL-C), a simple adaptation of the SPP model where instead of diffusing randomly, a cell moves in a directed fashion away from existing cell-cell contacts. Specifically,

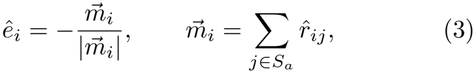

where 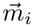 is the summed direction of cell *i*’s adhesive contacts, so the cell crawls opposite its net contact direction with fixed magnitude *f*_0_. (See Fig. 3 (a) (center) for a diagram). We show in Fig. 3 (b) (center column) that this model exhibits three distinct behaviors. For large off rates, the tissue falls apart into a loose collection of unmoving cells with no contacts. With increasing crawling speed and decreasing unbinding, an untensioned cluster phase emerges and grows. With enough activity, the cluster grows and stretches, spanning the box and forming the tensioned fluid phase. For large values of *v*_0_, CIL-C generates a uniform tensioned fluid that has a large diffusion coefficient, persistent tension and is more uniformly dispersed than random disks. In this state, the diffusion coefficient is set by *k*_off_. Additionally, examining the MSD reveals diffusive slopes are possible within the tension fluid regime, see SI. In this way, cells move in a locally directed manner, away from their neighbors, but globally undergo fluid-like diffusion, so a diffusive meansquared displacement alone cannot distinguish CIL from random motility.

Mesenchymal cells are also known for extending protrusions towards other cells. Therefore, we develop a second model based on binding new neighbors with a rate *k*_on_ and actively contracting or ratcheting that new cell-cell interaction with a rate *k*_*r*_, which we term CIL-P. (See Fig. 3 (a) (right) for a diagram). Candidate neighbors are defined using a Voronoi diagram, and to ensure cells move away from existing contacts we only make new interactions in regions where there is a large empty solid angle between existing adhesive contacts (see Methods for details).

We show in Fig. 3 that CIL-P generates a similar set of behaviors. Here we present data for *k*_on_ = 10^−3^ with data for *k*_on_ = 10^−4^ and *k*_on_ = 10^−5^ in the SI. With low activity and high unbinding, cells pull each other into untensioned clumps. However, with more activity and slower unbinding, a system spanning network supporting tension emerges. Diffusion is again controlled by the unbinding rate but also the ratchet rate, where increased unbinding increases the diffusion coefficient, as long as it does not deconstruct the network faster than new contacts are made. Taken together, CIL-P also is able to generate a tensioned diffusive fluid with a rearranging network.

### PSM cells show CIL-like behavior at the cell and tissue scales

Do PSM cells exhibit the CIL-like behavior our models require? To test for such behavior, we analyze the spreading of posterior PSM explants on fibronectin-coated substrates (see Methods), where CIL-like behavior should be apparent at two scales: in collisions between pairs of cells, and in the collective spreading of the explant.

At the cell scale, the mesenchymal nature of the tissue causes single cells to break away from the main cluster during spreading. These escaped cells either collide with other escaped cells (Fig. 4 (a)) or migrate without collisions (Fig. 4 (b)). We tracked *n* = 28 collision events from *n* = 7 multi-cluster explant experiments, in which two cells contact one another and then separate, with 30-minute windows before and after contact. We also identified *n* = 49 solo cell tracks, from which we extract independent 60-minute intervals for comparison to collisions, resulting in 302 non-interacting cell track pairs. In Fig. 4 (c), we align the pre-collision tracks along the *x*-axis with the collision at the origin, and color each post-collision track by its deflection away from the *x*-axis. We apply the same analysis to solo cells over two consecutive 30-minute windows in Fig. 4 (d), which provides a baseline for cell motion without cell-cell interactions. To compare the two populations, we compute the scattering angle *χ* between the relative velocities of the two cells before and after the collision (Fig. 4 (e)), where *χ* = 0° indicates no effect of the collision and *χ* = 180° indicates perfect scattering. Solo cells, paired as if they had collided, have a median scattering angle of 34° (95% CI 29–43°), consistent with non-interacting active Brownian particles with the observed single-track persistence time (see SI). In contrast, colliding cells have a median scattering angle of 99° (95% CI 71–130°) (Fig. 4 (f)) and the two distributions differ significantly (Kolmogorov–Smirnov *D* = 0.49, *p <* 10^−4^; see Methods). Thus, colliding PSM cells turn away from each other far more than expected for persistent random walkers, directly demonstrating CIL-like behavior between pairs of cells.

**FIG 4.**
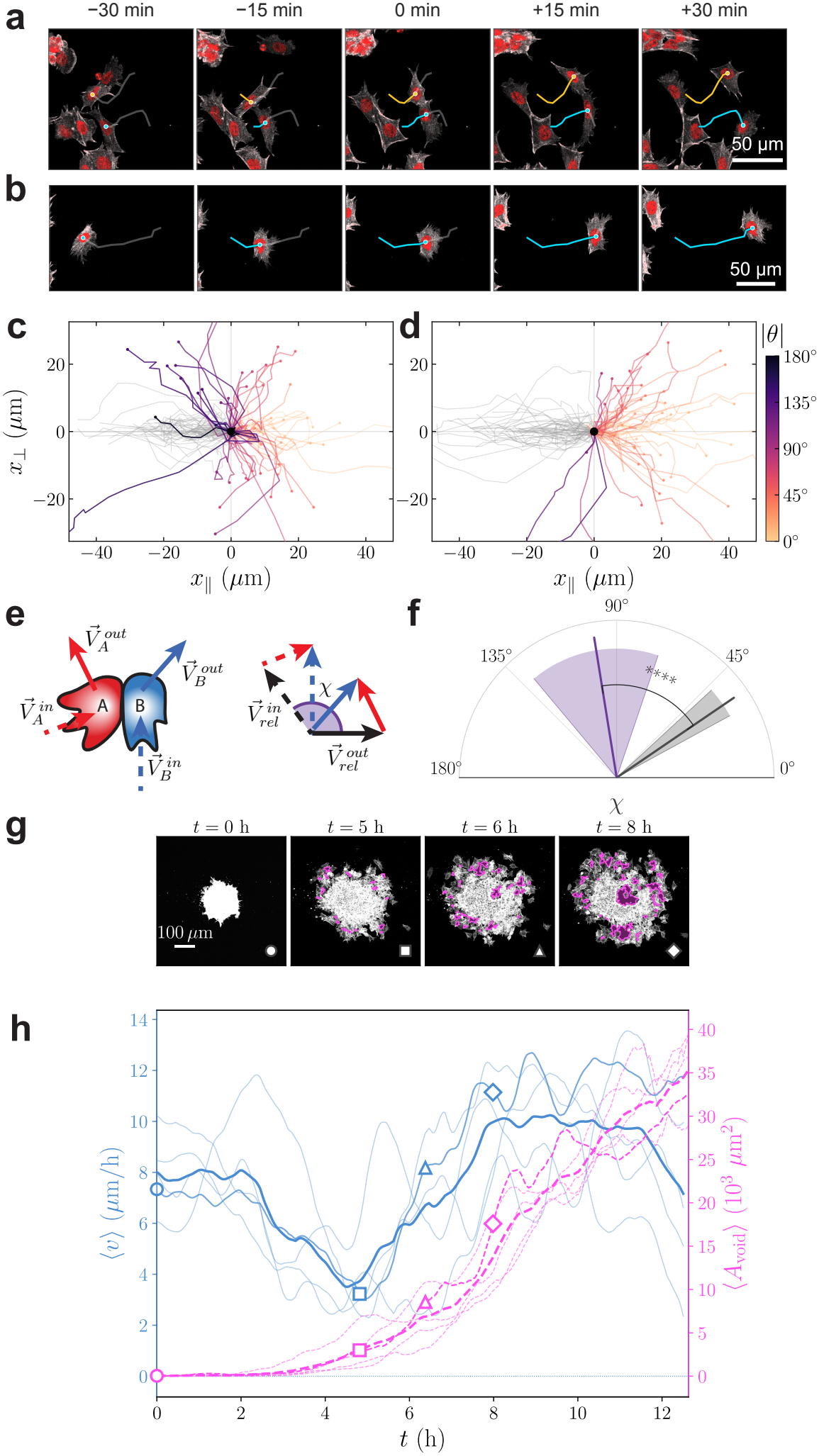
Explant spreading reveals cell-level and tissue-level CIL-like behavior. a) Example of a cell collision tracked over 30-minute time windows before and after contact. The cell bodies are labeled in white and the nuclei are labeled in red. Blue and orange tracks indicate where the cells have been, grey outlines indicate the track in the future. b) Example of a single cell moving with no interactions. c) Cell tracks from collision events are aligned so that the pre-collision track is along the x-axis from left to right and the contact event at the origin. Cell tracks after collision are colored by their deflection angle away from the x-axis, increasing from yellow to purple. d) Cell tracks for single cells that do not collide with another cell over two 30-minute windows. The first 30-minute windows are aligned along the x-axis from left to right (grey) to quantify the amount of turning in the following 30-minute window (yellow to purple). e) A diagram of two cells colliding and separating with their scattering angle calculated as the angle between their relative velocities before and after the collision. If the collision has no effect, the angle is 0 degrees. If the cells perfectly scatter the angle is 180 degrees. f) The median scattering angle for colliding cells (purple line, *n* = 28 events) and solo cells paired as if they had collided (grey line, *n* = 302 pairs from 49 cells) with their 95% confidence intervals (shaded fans). The two distributions differ significantly (Kolmogorov–Smirnov test, *p <* 10^−4^). g) An example of explant spreading. Pink areas indicate extracted void regions (see SI for higher-resolution images). h) Tissue-scale response of five explants, via quantification of their spreading. Blue (left y-axis) solid lines show the extracted velocity of explant spreading using a best-fit advection-diffusion model. Pink (right y-axis) dashed lines show the void area found via intensity thresholding. Thin lines indicate individual explant experiments. Thick lines indicate the average over 5 explants. Markers correspond to the images shown in g).

Does CIL-like behavior persist when cells are surrounded by neighbors? The spreading of cell aggregates has been extensively modeled as an active wetting problem, in which the balance of cell-substrate traction and cell-cell adhesion determines whether an aggregate spreads, accelerates, or dewets [58–60]. Of course, active wetting is consistent with CIL-like behavior, generating stress at the cluster edge driving wetting outwards. As a further test, we further ask what happens once the aggregate loses cohesion. Regardless of the details of traction or adhesion, CIL-like behavior predicts a distinct collective response once gaps open: as the outermost layer of cells moves outward and separates from the cells behind it, the second layer gains new empty space on its outward radial side. Via CIL, these cells should repolarize toward the new space, producing an accelerated and correlated outward motion of the tissue. If enough cells are activated to move radially outward, additional holes may open in the interior. As a tissue-scale assay of CIL, we therefore measure the radial velocity of the explant as holes first open.

We present an example spreading experiment in Fig. 4 (g). Initially, the explant spreads outward as a single disk. By 5 hours, single cells leave the cluster, by 6 hours holes open in the interior, and by 8 hours there is often a substantial gap in the middle of the interior. In a mathematical model where particles are simply diffusing, gaps will never open up in the interior; gaps generically require outward radial advection, where some of the particles/cells move outward faster than others. To quantify whether this advection is occurring and analyze its temporal variation, we fit each spreading experiment to both a diffusion and an advection-diffusion model. Indeed, we find that the diffusion-only model has nearly three times the mean squared error, ⟨MSE_*d*_⟩*/* ⟨MSE_*d*+*a*_⟩ = 2.7 *±* 0.2, so an advective model is a much better fit; explants do spread with an outward radial advection and are not purely diffusive.

The time dependence of the advective velocity helps to distinguish between two possible mechanisms: wetting only from the initial boundary or additional CIL-like behavior generated in the bulk. In Fig. 4 (h) we show a robust temporal signature averaged over five explants: the explant initially spreads with an advective velocity of ~8 *µ*m/hr. After approximately 5 hours, the velocity drops to a minimum of ~3 *µ*m/h as the first layer of cells leaves the cluster (square). Interior voids then open near the departing cells, coinciding with a sharp increase in outward velocity (triangle), before larger holes opens and the velocity plateaus again near ~8 *µ*m/h (diamond). To quantify the opening of holes, we measure the area of internal voids by intensity thresholding (Fig. 4 (h), pink). No voids are apparent before 5 hours, small voids appear between the escaped cells and the new cluster edge as cells break away, and the void area increases sharply as large central voids form near 8 hours.

If the mechanism was purely wetting at the boundaries, one might generally expect a constant outward velocity or a slowing down as the cohesion of the bulk tissue resists outward motion. In contrast, CIL-like behavior would suggest that the cells in the bulk are activated to move outward and increase the advective velocity precisely when voids start to open up and expose cells to additional free surface. Therefore, our observation of outward acceleration concomitant with void formation is supportive of a CIL-like mechanism.

Other cell-scale mechanisms are also likely contributing to void formation: the exact shape of the holes and the temporal evolution of the advective velocity are likely governed by a competition between CIL-generated stresses and cell-cell unbinding, analogous to the competition between *k*_*r*_ and *k*_off_ in our models. Although a detailed analysis is beyond the scope of the current work, explant spreading may prove a useful assay to constrain model parameters and feedbacks in the future.

Taken together, our observation of CIL-like behavior in pairs of cells and outward advection in cell aggregates confirm that cells in chick PSM do actively move away from neighbors, as predicted by our biophysical model.

### Continuum equations connect cell-scale features to emergent mechanics

Due to many-bodied interactions, spatial correlations, and the competition between contraction and relaxation, one might expect that a continuum theory to predict the behavior of *D, T* and 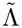 would present a significant challenge. Nevertheless, we find that mean-field assumptions generate a simple set of continuum equations, defined in terms of cell-scale parameters of the model, that are remarkably predictive of our simulation data for the CIL-P model. The crawling CIL-C model is more complex, as cell-cell binding driven by crawling-induced collisions introduces substantial spatial correlations. Even so, we find a similar scaling collapse to CIL-P presented in the SI.

We first consider the behavior of a single average spring representing an adhesive cell-cell contact. We assume that the binding, unbinding and contraction of the rest length of the spring is almost entirely controlled by simple two-state kinetics [61], and we ignore springs that form on contact. Then the time average contraction ⟨*c*⟩ of the spring rest length is

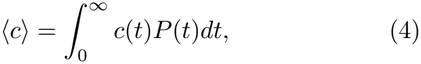

where *c*(*t*) = *k*_*r*_*σ*_*ij*_*t* linearly contracts the rest length from its starting length *l*_0_ until reaching the minimum contact distance *σ*_*ij*_, and *P*(*t*) is the model’s exponential unbinding probability. Solving this integral (see SI), we find

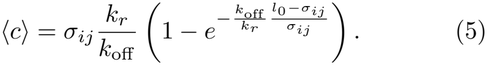

⟨*c*⟩ initially increases as *σ*_*ij*_*k*_*r*_*/k*_off_ until it saturates at *l*_0_ − *σ*_*ij*_, with the crossover at *k*_*r*_*/k*_off_ ~ (*l*_0_ − *σ*_*ij*_)*/σ*_*ij*_. Assuming a uniform distribution of cells, we have 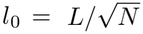, where *L* is the box size and *N* is the number of cells.

With the average contraction of the rest length, we can predict both *D* and *T*. First, consider the diffusion coefficient. If all of the contraction of a spring at some point is converted into motion via relaxation, for example by release of a neighboring spring, each binding event should displace a cell by *c* in a random direction. We should then expect *D* to scale as a random walk with step size ~ ⟨*c*⟩ taken at the flux of binding events, i.e. *D* ~ ⟨*c*⟩ ^2^*J*_*b*_ where *J*_*b*_ is the binding flux. In steady-state, assuming two-state kinetics leads to a simple equation for the binding flux: *J*_*b*_ = (*k*_on_*k*_off_)*/*(*k*_on_ + *k*_off_).

Similarly, if all the rest length contraction is converted into tension, as the cell is restrained by neigh-boring springs, we should expect the tension on each adhesion to scale with the spring force *k*⟨*c*⟩ times the separation *l*_0_, i.e. *T* ~ *k*⟨*c*⟩*l*_0_. The number of adhesions carrying this tension is set by the fraction of the total springs that are ‘on’, *f*_*b*_, as *N*_*b*_ = *Nz*_*max*_*f*_*b*_*/*2, where *z*_*max*_ is the maximum number of adhesions per cell. In steady-state, two-state kinetics results in a fraction bound of *f*_*b*_ = (*k*_on_)*/*(*k*_on_ + *k*_off_), which matches the measured number of adhesions to within 0.1 decades for *k*_on_ = 10^−3^ (see SI), so the total load carried by the network, *N*_*b*_*T* ∝ *f*_*b*_*k*⟨*c*⟩, follows the same two-state kinetics.

In Fig. 5 (a) and (b) we plot *D/J*_*b*_ and *T* vs *k*_*r*_*/k*_off_ respectively with the continuum results as dashed black lines. We find remarkable agreement between the simulation data and our proposed continuum model,

**FIG 5.**
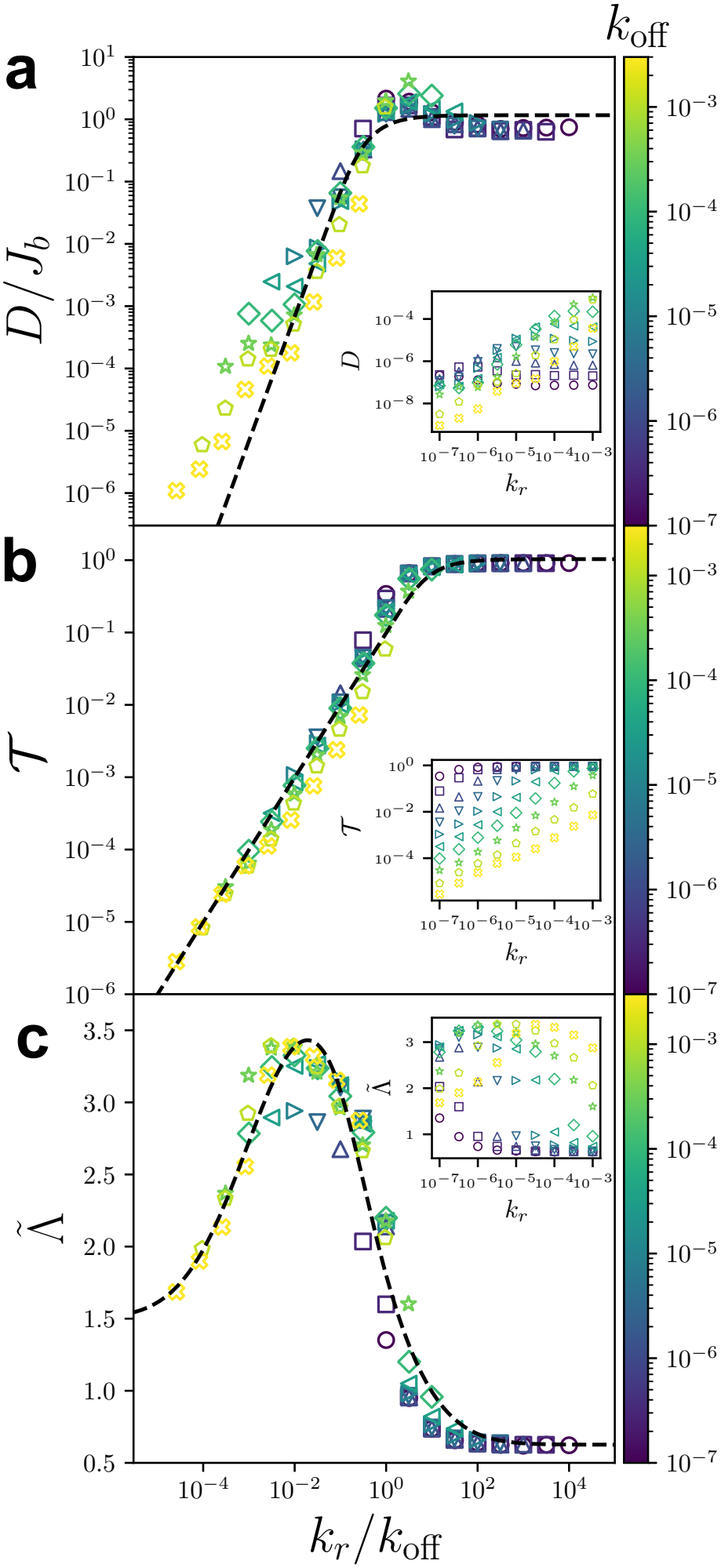
Continuum equations control the tensioned fluid state for CIL-P. (a) The diffusion coefficient relative to the binding flux *J*_*b*_, (b) the tension per adhesion *T* and (c) the normalized cell-scale texture 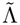 vs the ratchet rate relative to the off rate *k*_*r*_*/k*_off_. The black dashed lines in (a) and (b) indicate Eq. 6 with ⟨*c*⟩ from Eq. 5, where in (b) the crossover is at *k*_*r*_*/k*_off_ *≈* 10, and the black dashed line in (c) indicates Eq. 7. Insets present unscaled *D, T* and 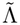 vs *k*_*r*_ respectively. Color indicates increasing off rate *k*_off_ from purple to yellow.

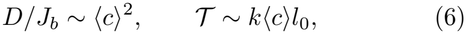

across 8 orders of magnitude of *k*_*r*_*/k*_off_ for the tension. The tension follows *T* ~ *k* ⟨*c*⟩ *l*_0_ in both limits, growing linearly with *k*_*r*_*/k*_off_ and saturating at the fully contracted spring, although the crossover between the two occurs at larger *k*_*r*_*/k*_off_ than for ⟨*c*⟩ alone (see SI). We present similar agreement for two further orders of magnitude in *k*_on_ in the SI. This collapse demonstrates that there is a steady-state flow of energy that is injected via the spring rest length contractions that governs both tension in the springs and motion of the cells as they relax this tension. Moreover, all the inputs for these continuum equations that predict the emergent dynamics are cell-scale features such as binding rates and distances between cells that can in principle be measured in future experiments.

The same two-state kinetics describe the crawling CIL-C model once crawling sets the on rate, *k*_on_ ∝ *v*_0_. The distance a released cell crawls before a new adhesion catches it then plays the role of *c* in the diffusion, the crawling force sets the tension, *T* ~ *γv*_0_ ⟨*σ*_*ij*_⟩, and the fraction bound, *D* and *T* collapse onto the two-state predictions, although less cleanly than for CIL-P (see SI).

Following this approach, we also find in Fig. 5 (c) that 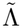 is well collapsed when plotted against *k*_*r*_*/k*_off_. 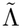 follows a similar behavior to Eq. 5, except that at low values of *k*_*r*_*/k*_off_ it saturates at whatever structure was imposed by the initial conditions, in our case 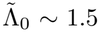, and there is also a saturating uniform tensioned fluid structure 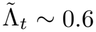 at high *k*_*r*_*/k*_off_, as the geometry prevents additional new contacts. Noticeably, there is a large peak in between these two limits as the tissue goes through a re-entrant clustering phase, with a maximum clustered structure 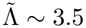. As an ansatz, we propose the correction,

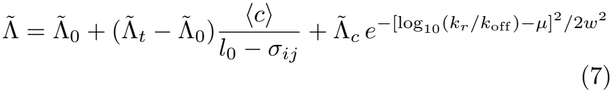

where the second term carries the texture from 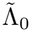 to 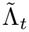 as the contraction saturates, and the last term is a log-normal peak of height 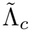 centered at *k*_*r*_*/k*_off_ = 10^*µ*^ with width *w* in decades. Fixing 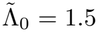 and 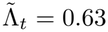, the measured plateau, we find the best fit 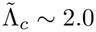, *µ* ~ −1.6 and *w* ~ 1.4, shown by the dashed line in Fig. 5 (c). A detailed justification for our texture ansatz must describe the emergence of clusters and likely requires a microscopic many-body theory, which we leave for future work.

## Discussion

While intercellular tensions are known to play an important role in confluent tissues, their contribution to the mechanics of non-confluent tissues remains poorly understood. Here, we show that the sparse cellular network of the avian PSM behaves as a unique material we term a fluid under tension, where at short time scales the cellular network retracts under tension in response to perturbations, while at longer timescales cells exhibit fluid-like exchange of neighbors and diffusive motion. In this sense, the tensioned fluid resembles a viscoelastic material, responding elastically on short timescales and relaxing stress through adhesion turnover on longer ones. Unlike a passive viscoelastic fluid, however, it relaxes perturbations while remaining under tension, because CIL continually regenerates the adhesion network. This behavior also differs from the jamming transition that describes the more densely packed zebrafish PSM [7]. Although one might expect a clustering instability in a porous rearranging material under tension, the local cellular structure is instead *more uniformly dispersed* than randomly placed spheres.

We explain this using a simple mesenchymal cell-cell interaction model, and demonstrate that fluid under tension properties are not observed unless the activity is directed away from neighbors. This activity is reminiscent of CIL and combats the tension instabilities that would otherwise collapse a porous cellular network. Moreover, we develop a set of continuum kinetic-based equations and effectively predict how cell-scale features control diffusivity, tension and the structural texture. This immediately suggests new experiments to measure these cell-scale features, such as adhesive cell-cell contact on- and off-rates and effective ratcheting rates, in order to connect them to regulation and disregulation of PSM architecture and function. For example, in the avian PSM there is a strong gradient in cell and tissue properties along the anterior-posterior axis [9]. Experimentally quantifying and perturbing cell-cell binding and unbinding rates and contractility along that axis would allow our model to provide testable predictions for tension and structure gradients, as well as tissue-scale response to perturbations.

Our work complements a previous theory/simulation study that explored how reorientation time and crawling activity organize a tissue [62]. That manuscript reports a region of phase space that does behave like a tensioned fluid, but the texture is clustered, similar to our SPP model and unlike the uniformly dispersed CIL models that match the PSM. The absolute tensions in our CIL models may also be higher, as the cell-cell potential used in [62] is similar to a sticky particle, which does not permit large tensions in the contacts before they break, instead of the hysteretic potential employed here. Future numerical work should use quantitative measures like the viscoelastic moduli to compare and tease apart how different forms of activity and interaction potential alter the emergent mechanical response.

Of course, the major prediction of this work is that some cellular behavior like CIL is operating in stellate mesenchymal tissues. While CIL has long been studied in single and collective cell migration [63–65], less attention has been paid to how CIL might organize entire tissues [52]. We use the term CIL-like because our models capture a behavior, cells moving or protruding away from existing contacts, rather than a specific molecular mechanism, and the PSM may produce this behavior through pathways different from those characterized for CIL in other cell types. Our explant experiments provide initial evidence for CIL-like behavior in the PSM at both scales: colliding cells turn away from each other far more than non-interacting cells, and spreading explants accelerate outward as interior voids open. As this initial work is on fibronectin-coated substrates outside the embryo, it would be interesting in the future to develop new cell- and tissue-scale techniques to search for CIL behaviors *in vivo*, in the PSM and other mesenchymal tissues.

Focusing on the cellular network of the PSM also highlights potential connections to molecular-scale networks that undergo reorganization [66]. Concurrently to this work, a similar model of a spring network was proposed to explain actin reorganization [67]. Similar to the effects of CIL at the cellular scale, that work proposes such networks add new connections in directions of lower density. The fact that both teams of researchers independently identified similar mechanisms suggests that the tensioned fluid phase could describe a large class of materials over different length scales across biology or materials design. While both CIL-C and CIL-P models generate similar phases, the models are not identical; CIL-P is selflimiting in a way CIL-C is not. In the pulling model, a cell can only reach out and bind its nearest neighbors, putting a limit to how fast they can be pulled in. In the crawling model, the crawling force can be continually increased, and at sufficient magnitude the active force pushes the cells beyond their nearest neighbors. Future work should further delineate these two models and identify features that might distinguish them in experiments. This also paves the path for future modeling and experimental work that investigates how key biological feedbacks alter cell-scale parameters. For example, cell-cell unbinding rates likely depend on the tension across the cell-cell interaction, and CIL dynamics and reorientations are likely impacted by molecular signals such as FGF, an established regulator of PSM fluidity [12]. In addition, previous work on cartilage [56] suggests that an interstitial soft ECM network, such as hyaluronic acid in the PSM, will play an important role in modulating the rigidity and fluidity of the composite tissue. We expect such soft ECM will provide a restoring force, similar to CIL, that opposes the tension on short and intermediate timescales, although the ECM must be able to reconfigure on longer timescales to allow neighbor exchange, and therefore we still expect CIL to be necessary to prevent clustering on longer timescales. The ECM may also connect our results to models in which axis elongation is driven by extracellular volume expansion [9]. A swelling hyaluronic acid gel can be under pressure while the embedded cellular network remains under tension, so that expansion pushes against a tensioned, rearranging backbone of cells rather than only against the tissue boundaries. Testing this composite picture will require models that couple the cell network to a swelling gel, which we leave for future work. In short, our new characterization of the cell-scale features that are important for fluid-under-tension behavior in sparse cellular networks drives testable new hypotheses for how biomolecular players control tissue form and function during mesenchymal morphogenesis.

## METHODS

### Animals

Fertilized chicken eggs were obtained from the Roslin Institute (Edinburgh, UK). Three transgenic lines were used in this study: (i) the ACTN line, which exhibits ubiquitous expression of Lifeact-Venus and nuclear H2B-mCherry [68]; (ii) the Roslin Green line, which shows ubiquitous cytoplasmic eGFP expression [69]; and (iii) the Chameleon line, which carries a ubiquitous nuclear blueFP reporter that is replaced by eYFP, tdTomato, or mCerulean following Cre-mediated recombination [68]. Embryos were incubated at 37° C for 42–44 hours until reaching stage HH9–10.

### Time-lapse microscopy

For time-lapse imaging, HH10 transgenic embryos expressing H2B-mCherry were harvested using the filter paper technique described by Chapman et al. [70] and cultured ventral side up at 37° C for 2–3 hours. Embryos were screened for fluorescence intensity and transferred ventral side down onto imaging dishes coated with a thin albumin/agarose layer. All prepared embryos were then placed in the microscope incubator and maintained at 37° C. For cell-tracking experiments, nuclear labeled embryos were imaged every 2 minutes for 2 hours, acquiring 10–12 z-planes at 1 *µ*m intervals.

Nuclear labeled image tracking was performed using Ultrack [32]. Foreground and background regions were automatically identified using Ultrack’s intensity-based foreground detection, and nuclei were segmented across z-stacks using standard Ultrack segmentation and watershed refinement. Object linking across time points employed Ultrack’s global optimization tracking framework, which minimizes spatial displacement and intensity-based costs to reconstruct continuous cell trajectories. Tracking employed Ultrack’s global association solver. The configuration parameters were tuned as follows: tracking window size of 20 frames, overlap of 15 frames, a maximum linking distance of 4.0 *µ*m, a z-score threshold of 1.2, appearance/disappearance weights of −0.20, division weight of −4.0, and inclusion and dismissal weights of 1.0 and −1.0, respectively.

### Laser ablation

Transgenic embryos expressing cytoplasmic GFP (Roslin Institute, UK) were harvested at the 10–12-somite stage and cultured in Early Chick (EC) conditions [70]. After 1 hour of incubation, embryos were mounted ventral-side down in direct contact with glass-bottom dishes coated with embryo culture medium. Embryos were then transferred to the heated (37° C) stage of an inverted Zeiss LSM880 confocal microscope. This setup prevented embryos from drying and kept them in a stable position during imaging and ablation. Laser ablations were performed as described by Marshall et al. [71]. To make small, localized cuts without cavitation, ablations were performed at 40x magnification and targeted to individual cell protrusions. Multiple ablations were performed along the mid-anteroposterior region of each PSM column, for a total of 25 ablations across 7 embryos. A z-stack spanning the ablation site was acquired immediately after ablation to verify that ablations were confined to the PSM, approximately 20–30 *µ*m from the ventral surface, with the overlying endoderm intact (see SI). The tissue relaxation following laser ablation was captured using a PIV method previously reported in Saadaoui et al. [35], using a PIV interrogation window of 8 *µ*m to capture cellular-scale movements.

### Explant preparation and imaging

Posterior PSM (P-PSM) explants were dissected from Lifeact-Venus and nuclear H2B-mCherry transgenic chicken embryos (Roslin Institute, Edinburgh, UK) at the 10–12-somite stage. Explants were incubated in culture medium containing DMEM/F12 (Gibco) supplemented with 10% FBS (Gibco) and 1% Pen/Strep (Gibco) at 37 °C and 5% CO_2_. After 30 min, explants were transferred to fibronectin-coated glass-bottom dishes, followed by an additional 2–3-hour incubation. Time-lapse imaging was performed using a Fluoview FV4000 confocal laser scanning microscope (Evident/Olympus), and images were acquired every 2–3 min for 450–500 cycles. We analyzed *n* = 5 single-cluster explants for collective spreading and *n* = 7 multi-cluster experiments, in which several clusters were plated together to increase the likelihood of observing single-cell collisions.

### Explant single-cell tracking

In the *n* = 7 multi-cluster experiments, cells break away from the clusters and crawl freely on the substrate, and we track these free cells by hand. We built an annotation tool in napari [72] that overlays the Lifeact-Venus and H2B-mCherry maximum-intensity projections, and we follow each cell by clicking the center of its nucleus in every frame. We record two classes of tracks. Solo tracks follow a free cell that does not interact with any other cell; we track every free cell that crawls for at least two consecutive 30-minute windows without an interaction, giving 49 solo tracks. Collision tracks follow two free cells that collide and subsequently separate, identified by eye. For each collision, we track both cells through the approach, contact, and separation, and we mark a single contact time *t*_*c*_ at the approximate midpoint of the interaction, since cells often remain in contact for several frames. We tracked *n* = 28 collision events across the seven experiments. Two annotators (A.T.G. and G.S.P.H.) tracked cells in the same movies, and for cells tracked by both we keep one annotator’s track.

### Collision analysis

For each cell, we measure the velocity before and after contact by a least-squares linear fit of position versus time over the windows [*t*_*c*_ − *w, t*_*c*_] and [*t*_*c*_, *t*_*c*_ + *w*], with *w* = 30 min. The scattering angle *χ* is the unsigned angle between the relative velocity **v**_rel_ = **v**_*A*_ − **v**_*B*_ of the two cells before contact and after contact, so that each collision gives one value of *χ* ∈ [0°, 180°]. For the null, we pair solo cells as if they had collided, placing a pseudo-contact at every frame for which both cells are tracked over both windows, and measure *χ* exactly as for a collision. Pairs that share less than 2*w* of tracked time are excluded, leaving 302 pseudo-pairs from 49 cells.

Because pseudo-contacts within one pair overlap in time and each cell appears in many pairs, we reduce each pseudo-pair to its median *χ* and estimate the 95% CI of the median with a pigeonhole bootstrap that resamples cells [73]; for collisions, we bootstrap over events. For the Kolmogorov–Smirnov test, we thin the null so that pseudo-contacts within each pair are at least 2*w* apart and report the 90th percentile of *p* over 500 random thinnings.

### Explant spreading analysis

We analyze the maximum-intensity projections of *n* = 5 explants imaged in a single acquisition (0.621 *µ*m per pixel, 2.53 min per frame), until the spreading cluster reached the edges of the field of view (~13 hours). For each frame, we compute the intensity-weighted center of mass and average the pixel intensity in annuli of width Δ*r* = 1.24 *µ*m to obtain the radial density profile *ρ*(*r, t*). To remove the background, we subtract from each profile a single value, the median of *ρ* over 0.80–0.98 of the largest radius covered by pixels in every frame, and clip the result at zero. We subtract one value rather than thresholding each pixel because a per-pixel threshold preferentially removes the dim pixels in gaps between cells, whose number grows as the explant spreads, and would bias the interior density in time. Each profile is normalized to unit mass, ∫ *ρ dA* = 1.

We model the spreading explant as an axisymmetric density evolving under diffusion and a radially outward advective velocity *v*,

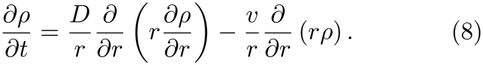

For *v* = 0, a disk of density spreads into a wider, lower-density disk whose maximum remains at the origin. For *v >* 0, the radial velocity field has divergence *v/r*, which drains the center and builds density at the edge. We solve Eq. (8) with a mass-conserving finite-volume scheme on the radial grid of the data, extended to 955 *µ*m so that less than 10^−5^ of the model mass reaches the outer boundary, with upwinded advection, zero-flux boundaries, and explicit Euler time steps of Δ*t* = 1–6 seconds, below the diffusive stability limit Δ*r*^2^*/*4*D*. For each explant, we propagate the measured profile at *t* = 0 and fit constant *D* and *v* by minimizing the squared error between the model and data profiles, MSE(*t*) = ∫ (*ρ*_data_ − *ρ*_model_)^2^ *dA*, summed over all frames, starting from a coarse grid search followed by Nelder– Mead refinement. We fit a diffusion-only model (*v* = 0) with the same initial profile, solver, and objective, so that the two fits differ only by the advective term, and compare them by the ratio of their time-averaged errors, ⟨MSE_*d*_⟩ */* ⟨MSE_*d*+*a*_⟩, reported as the mean ± standard error over explants.

To resolve how the advective velocity changes in time, we fit *v*(*t*) sequentially along a single model trajectory, holding *D* fixed at each explant’s constant-velocity value so that all of the time dependence is carried by *v*. At frame *i*, the initial condition is the model’s own prediction from the previous step, not the measured profile. We fit *v*_*i*_ by bounded Brent minimization of the summed error over the next 8 frames (20 min), advance the model by one frame under *v*_*i*_, and repeat. We average *v*(*t*) over explants frame by frame and display it with a centered 0.75 hour running mean. The timing of the minimum and recovery of ⟨*v*⟩ is insensitive to the fitting window (4 to 16 frames), to the choice of center, and to using the H2B-mCherry nuclear channel in place of Lifeact. The magnitude of *v*(*t*) depends on the fixed *D*, which trades off against *v*: halving *D* raises ⟨*v*⟩ by 1–2 *µ*m/h and doubling *D* lowers it by about 2 *µ*m/h, but the overall trends remain the same.

To quantify the opening of voids, we segment each frame of the same movies used for the density profiles. Each frame is divided by its mean intensity, smoothed with a Gaussian filter of width 2 pixels, and thresholded in log_10_ intensity with Otsu’s method [74], recomputed for each frame. The explant envelope is the largest connected component after a morphological closing with a disk of radius 3 pixels and removal of objects smaller than 500 pixels, with enclosed regions filled. Voids are the regions inside the envelope that are not cell material in a separate mask closed with a disk of radius 1 pixel, so that small voids are not erased by the stronger closing used for the envelope. Islands of cells disconnected from the main body are not counted as void. To separate voids from the gaps between neighboring cells, we retain only voids larger than half the area of a 10 *µ*m-diameter cell (39 *µ*m^2^). The void area *A*_void_(*t*) therefore includes both interior voids and the gaps enclosed between the explant edge and departing cells. We average *A*_void_(*t*) over explants and display it in the same manner as *v*(*t*).

### Texture analysis

To quantify the tissue structure, using the same method for both the microscopy and simulations, we compute Λ, the variance over all window positions of the fill fraction *λ* in a square window of side *r*, on binarized images. For the experimental data, we use maximum-intensity z-projections of the nuclear-labeled movies, restricted to the PSM by fixed masks on either side of the neural tube, and divide the masks into non-overlapping 140 *µ*m × 140 *µ*m images. In the case of the experimental images, background noise in the image would artificially increase the uniformity. However, over-smoothing the image would also increase the uniformity. Therefore, we apply a minimal smoothing filter of a Gaussian blur with a width of a tenth of the average cell radius. Because the image brightness varies across the field of view and over time, we then normalize the local contrast over windows of ~5 cell diameters and binarize each image using Otsu’s method [74]. The observation window is set to the average cell diameter, *r* = 12*µ*m, and the texture of each image is normalized by that of RSA disks at the same fill fraction, rendered with the same pixel size and observation window. The averages and standard errors reported for the experimental data were taken over 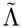 per image. For the simulation data, images were rendered as circles, without drawing adhesive springs as they have no extent in the simulations, and the observation window was set to the average diameter. See SI for further details.

To illustrate the range of possible textures, we generated several reference datasets beyond RSA. To create a highly uniform sample, we placed repulsive disks at a high packing fraction of *ϕ* = 0.84 and energy minimized the system using FIRE to an average force magnitude of |*F*| *<* 10^−14^. Then, we uniformly shrunk the disk sizes until the target packing fraction matching the experimental data of *ϕ* = 0.4 was reached. For clustered samples, we placed repulsive disks with an attractive shell at *ϕ* = 0.4 and energy minimized the system. The longer range the attraction, the larger the clusters. We find 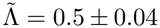 for the diluted jammed packings and 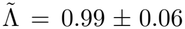 for the random discs by construction.

Our most clustered samples, where every disc is in the same cluster, achieve 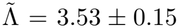, the approximate upper bound of clumpiness for our cell-scale texture metric.

### Numerical simulations

All simulations were conducted by integrating Langevin dynamics,

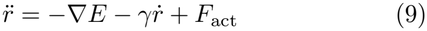

where *r* is the positions of the cells, *E* is the interaction energy, *γ* is a linear damping term and *F*_act_ is the active force of the particular model, with unit mass. The simulations were conducted in the overdamped limit, with no temperature fluctuations. The system was integrated using the BAOAB integration scheme with a step size Δ*t* = 0.1 [55] and an input damping of 10^3^. Because 10^3^ Δ*t* ≫ 1, the velocity is fully damped within each step and the scheme reduces to overdamped Euler integration, 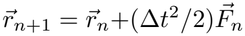, i.e. Eq. 9 in the overdamped limit with an effective damping *γ* = 2*/*Δ*t* = 20, independent of the input value. Throughout, *γ* refers to this effective damping, so that the free swim speed is *v*_0_ = *f*_0_*/γ*. Simulations were conducted with *N* = 256 cells under periodic boundary conditions and run for 10^9^ integration steps and with measurements reported after reaching steady-state and averaged over 10 replicates. Simulations were conducted in 2 spatial dimensions, as the tissue region studied is approximately only 5 cells thick along the dorso–ventral axis.

The interaction energy was modeled as our new hysteretic sticking potential as described in the main text. Cells with diameters *σ*_*i*_ do not interact at a distance greater than their repulsive contact distance, the average of their diameters is *σ*_*ij*_ = (*σ*_*i*_ + *σ*_*j*_)*/*2 and diameters were chosen from a random uniform distribution between 1 − 1.4 to suppress crystallization, since crystallization is not observed in experiments. This size ratio is standard for suppressing crystallization in two-dimensional packings [26], and the resulting polydispersity, the ratio of the standard deviation to the mean diameter, is ≈ 0.1. This value is comparable to the ≈ 0.17 we estimate for the PSM from the nuclear segmentations used for tracking, taking each nucleus’s diameter as that of a circle with its track-averaged area. Once two cells are closer than *σ*_*ij*_, they interact with a double-sided linear spring,

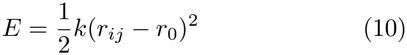

where *k* is the spring constant and *r*_0_ is the spring rest length. At contact, *r*_0_ is set to *σ*_*ij*_ to represent an unstressed cell-cell contact forming. Compression of the spring represents deformation of the cells coming closer together. Stretching of the spring represents the deformation of the cell as adhesive arms stretch out from the cell bodies as well as the forces at the intercellular junction and ECM-mediated forces. Compression and extension spring constants were equal in CIL-C and CIL-P, however; in the SPP model the compression spring constant was increased to be 100 times larger to prevent large overlaps for large values of *v*_0_. This was unnecessary for the CIL models as they reorient away from new contacts instantaneously causing very little repulsive overlap.

Unbinding events were performed kinetically at random to allow for the remodeling of the cellular network. Every *N*_steps_, every cell-cell contact was checked for unbinding by drawing a uniform random number *x* between 0 and 1 and the spring representing a cell-cell contact would be removed if the random number was *x < p*_off_. This results in an exponential probability of unbinding 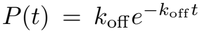 and an unbinding rate *k*_off_ ~ *p*_off_ */*(*N*_steps_Δ*t*). *N*_steps_ = 10^3^ was selected as a balanced for computational efficiency and accuracy of the kinetics.

For the SPP model, we chose to study the active Brownian particle (ABP) [53]. In the ABP the active force,

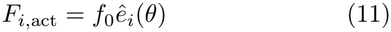

where *f*_0_ is the active force magnitude, resulting in a swim speed of a free particle of *v*_0_ = *f*_0_*/γ*, and 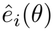 is the unit director of the activity of cell *i*. The unit vector evolves according to a stochastic Brownian process controlled by the rotational diffusion rate *D*_*r*_, which was set to the small value of *D*_*r*_ = 10^−4^ for all ABP simulations.

For the active crawling CIL-C model, the active force was deterministic, pointing away from a cell’s adhesive contacts with fixed magnitude *f*_0_,

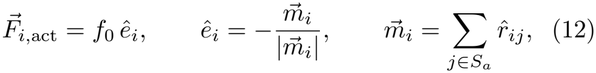

where *S*_*a*_ is the set of cell *i*’s adhesive contacts and 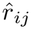 is the unit vector pointing from cell *i* to cell *j*, as in Eq. (3). A cell with no adhesive contacts has no active force. The reorientation of activity direction upon contact is instantaneous.

For the active pulling CIL-P model, there is no additional active force. Instead, the CIL-P activity is modeled using the interaction energy between cells by introducing new cell-cell contacts and pulling on them. Binding events were considered every *N*_steps_ at the same time as unbinding events. In order to carry out a stable active pulling model, cells need to identify two conditions: 1. Which of all the cells in the system are possible binding partners? and 2. What local conditions between two potential binding partners should be met?

First, we have found that the simplest and most efficient manner of identifying possible binding partners is using a Voronoi tesselation. Using a simple distance cutoff criteria appears intuitive, however, such a method leads to a feedback in which as a cluster begins to form, the list of potential binding partners will be biased towards the cells in that cluster, causing the cells to pull on the cluster and increasing its size until the system collapses into one large cluster. We found no set of parameters with a distance cutoff to be stable over time. Additionally, we motivate the Voronoi tesselation as representing that a cells ability to reach out and grab another cell is limited to its nearest neighbors. We used Voro++ to draw the Voronoi tesselation [75]. For computational efficiency, the Voronoi tesselation was only drawn every *N*_steps_ when all binding events were considered.

Second, in order to mimic a CIL type behavior, we require both potential binding cells to follow a set of local rules. The general principle is that for each cell to reach out a new arm and bind another cell, there needs to be enough cell surface available that is not already involved in an adhesive contact. To accomplish this, for each considered cell-cell pair, we calculate all the angles between each existing cell-cell contact by measuring the absolute angle of the cell-cell contact vector with atan2(y,x), sorting this list and then taking the difference in neighboring absolute angle. We then require that both cells have an angle that is larger than a cutoff *θ*_*m*_. Finally, for a new cell-cell adhesive contact to reach out between two neighboring cells that satisfy the *θ*_*m*_ criteria, these two open patches must point at each other. We check this by making sure the separation vector between the two cells is within the satisfying angles by adding it to the sorted absolute angle lists. For this study *θ*_*m*_ = 2 rad. This affects both the maximum number of possible cell-cell contacts and the fraction bound curve. With *θ*_*m*_ = 0, the maximum number of neighbors converges to the Voronoi neighbors of *z* ~ 6. *θ*_*m*_ = 2 results in a more biologically relevant *z* ~ 4.5, but we leave a precise exploration of this parameter for future work.

After forming a new cell-cell contact through the binding mechanism, the initial rest length is set to *r*_0_ = |*r*_*ij*_|. The cell-cell contact is then pulled in by decreasing this rest length by Δ*r*_0_ = *k*_*r*_*σ*_*ij*_Δ*t* where *k*_*r*_ is the ratchet rate. *r*_0_ is decreased by Δ*r*_0_ every integration step. To respect contact distance, the ratcheting ends when *r*_0_ = *σ*_*ij*_ and is held constant.

### Numerical tension

In the numerical simulations, we define the tension as the negative virial pressure per cell-cell adhesion. The virial stress is computed using the virial stress tensor [55],

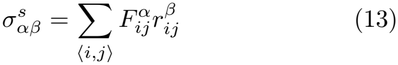

where *α, β* ∈ *{x, y}*, 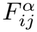 is the *α*-component of the force between cell *i* and *j* and 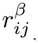 is the *β*-component of the separation vector between cell *i* and *j*. The virial stress contribution to the pressure in 2 spatial dimensions is then,

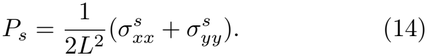

With *N*_*b*_ adhesions in the system, the tension is

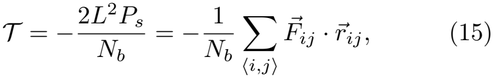

the average force times separation carried by an adhesion, which is positive when adhesions are stretched.

We pool both sums over the steady-state frames and replicates of each state point, and set *T* = 0 when no adhesions remain. The thermal contribution to the pressure from velocity fluctuations [55] and the active swim pressure of the SPP and CIL-C models [76, 77] are not carried by adhesions and are not included; in the CIL models both are below ~1% of the virial term in the tensioned fluid phase.

## Code Availability

The discrete element simulation code developed here is available at: https://github.com/agrigas115/hysteretic_sticking_plus_CIL.

## ACKNOWLEDGMENTS

AMongera thanks the members of the Multicellular Morphogenesis Lab for insightful discussions, and in particular Louis Hamilton for assistance with egg handling and incubation; Claudio Stern for generously sharing laboratory space, instruments, and reagents; the Centre for Cell and Molecular Dynamics for support with the *in vivo* imaging experiments, especially Alan Greig; and Dale Moulding at ICH for assistance with the laser ablation experiments. AMongera and GH acknowledge financial support from the Academy of Medical Sciences (SBF009\1014), the Human Frontier Science Program Organization (RGEC31/2024), the Royal Society (RG\R1\251273) and the Wellcome Trust (319420/Z/24/Z). EM acknowledges funding from Taighde Éireann-Research Ireland under grant agreement 24/PATH-S/12626. GLG acknowledges support from the Leverhulme Trust (RPG-2024-147). AMichaut thanks Francis Corson for sharing the PIV software he developed. MLM and ATG acknowledge support from grant number 2023-329572 from the Chan Zuckerberg Initiative DAF, an advised fund of Silicon Valley Community Foundation and Syracuse University’s Orange Grid research computing cluster. ATG thanks Rebecca Grigas for assistance with graphic design.

