## Supplemental Information for "Sparse mesenchymal cell networks as a fluid under tension"

The Supplemental Information includes additional information and analyses to support the results and conclusions in the main text. In Section 1, we present further information on the nuclei labeled tracking data and vary distance cutoffs in the displacement analysis. In Section 2, we confirm that the laser ablations target the PSM and leave the ventral surface of the embryo intact. In Section 3, we compare reference texture curves for Poisson points and randomly placed non-overlapping disks as a function of increasing density. Additionally, we consider different window sizes for analyzing the nuclei labeled data, show that a similar texture can be seen in the cytoplasmically labeled cells, and consider randomly placed non-overlapping ellipses for normalizing the texture data. In Section 4, we show that the scattering angle of non-colliding cells matches that of non-interacting active Brownian particles. In Section 5, we show the explant images of the main text with and without the extracted voids. In Section 6, we show example diffusive mean-squared displacement curves in the tensioned fluid phase for both CIL-C and CIL-P models, along with visualizations of the frozen, clustered and tensioned fluid phases. We also present the collapse of the CIL-P data as a function of binding rate $k_{\text{on}}$ and test the robustness of the tensioned fluid to cell division and death. In Section 7, we outline further analytical details on the tensioned fluid phase in both models, including the kinetic theory and collapse for the CIL-C model. Lastly, in Section 8, we provide captions for the supplemental videos.

### 1 Nuclei tracking

In the main text, we present a small patch of the nuclear tracking in the PSM. Here, in Fig. S1 (a) we show the entire tracking of the developing PSM for 2 hours for one of the two embryos that was imaged. Tracks are colored by time from blue to red. The center gap represents the masked notochord, flanked by two portions of PSM tissue. We do not plot segments where the cell was tracked for less than 10 minutes continuously and ignore these short tracks for downstream data analysis. In Fig. S1 (b), we plot the probability distribution of continuous track lengths extracted. The peak is at four minutes and decays exponentially. We successfully extracted 3,459 continuous nuclear tracks that are longer than 10 minutes, with an average length of 18 minutes long. The decrease in longer cellular tracks is the basis for the increasing standard error over time in the main text and why displacement metrics were cutoff after $\sim$25 minutes.

As further evidence of the diffusive nature of the PSM cell tracks when elongation drift is removed, we consider varying the cutoffs used to define neighbors. In Fig. S2 (a), we show how the mean-squared difference in neighbor distances remains approximately linear when varying the cutoff in initial neighbor distance $r$. We test both shorter and longer distances than the average cell diameter of 12 $\mu$m. Additionally, in Fig. S2 (b), using the same initial neighbor cutoff definition of 12 $\mu$m, we vary the cutoff in the distance to be classified as no longer neighbors $r$. As the cutoff is increased, the fraction of neighbors that rearranges decreases, but the result is robust up to the approximate average 5 $\mu$m distance displacement seen in Fig. S2 (a) after 20 minutes.

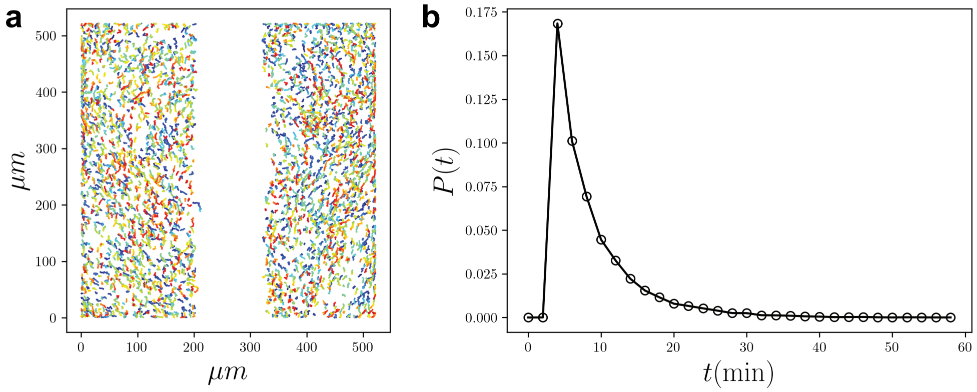


Figure S1: (a) Nuclear tracking of the developing PSM over 2 hours. Individual line segments represent continuous tracking of the same cell over time, colored from blue to red to indicate increasing time. (b) Probability distribution of the extracted nuclear cell track lengths in minutes.

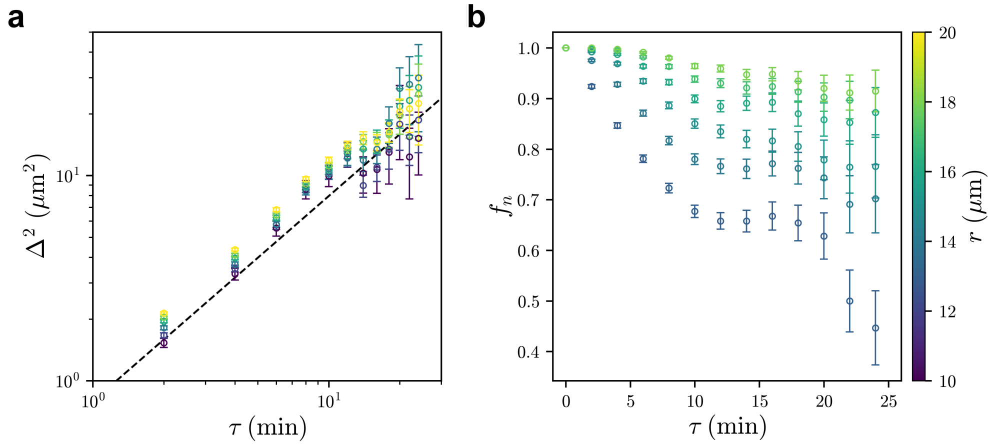


Figure S2: (a) The mean-squared distance displacement $\Delta^2$ vs lag-time $\tau$ for tracked nuclei that are initially neighbors $|r_{ij}|(t) < r$ . (b) The fraction of initial neighbors $f_n$ that remain neighbors vs lag-time $\tau$. Color indicates by various neighbor cutoff distances $r$ increasing from purple to yellow in $\mu$m. Error bars indicate the standard error.

#### 2 Laser ablation targeting

To confirm that the PSM was specifically targeted during ablation, we acquired a z-stack encompassing the ablation site immediately after the cut was made. In Fig. S3 (a), we present an example showing that the ventral surface of the embryo, including the endoderm, remains intact following ablation. In Fig. S3 (b), we quantify this by comparing the pixel intensity above the ablation site with that outside the column corresponding to the ablation site. The average intensities are nearly identical, indicating that the tissue above the ablation site remains largely unaffected. Furthermore, this analysis confirms that the ablations were consistently targeted to the PSM, at a depth of approximately 20–30 $\mu$m images from the ventral surface.

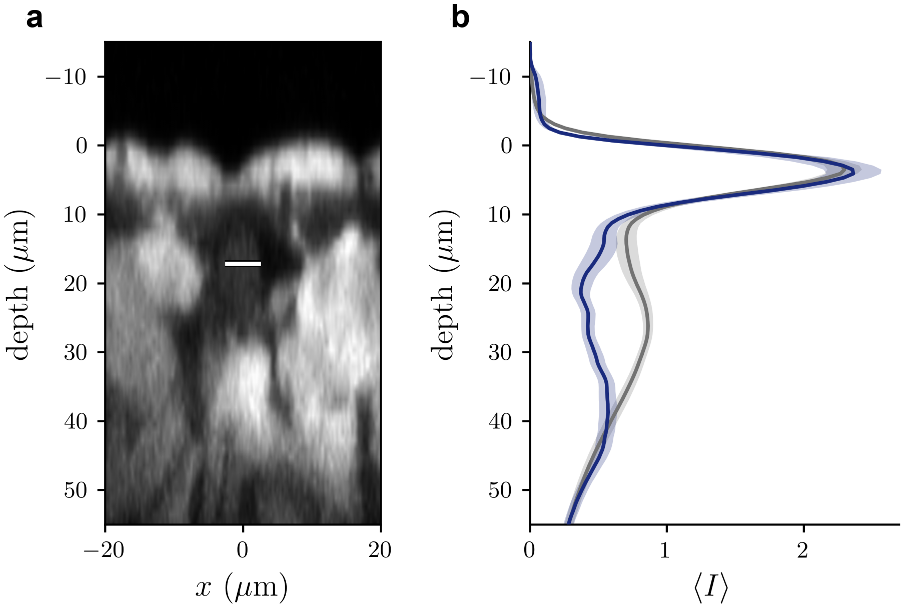


Figure S3: (a) XZ projection example of the PSM post ablation. The white bar indicates the targeted cut location. A gaussian filter was added to smooth the z-axis. (b) Quantification of GFP intensity along the ventral (top) to dorsal (bottom) axis of an embryo cross section. Using the ablation region, we define a column around the cut location (blue) and outside of the cut column (grey). The edge of the endoderm is aligned and the average intensity from the void region is subtracted. The intensity is further normalized by the average of the region outside of the cut column. Solid lines are the mean over n = 24 cuts in 7 embryos, and shaded bands are the standard error of the mean across cuts at each depth. The intensity starts at zero above the edge of the endoderm. Then the intensity peaks near 4 $\mu$m, indicating the endoderm. But inside and outside of the cut column show the same intensity within error, indicating the cut does not damage the surface. The cut region is about half as bright as the outside from 20–30 $\mu$m.

### 3 Cell-scale texture

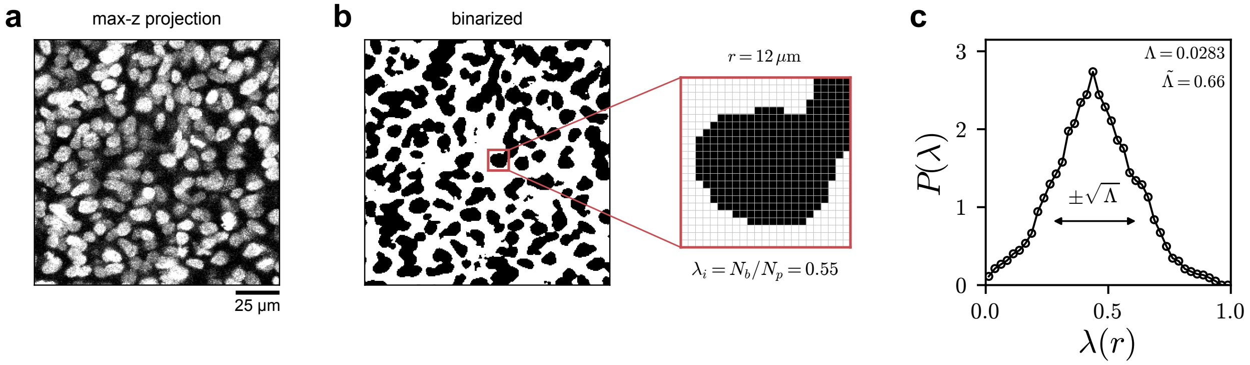


Figure S4: Calculating the normalized cell-scale texture $\tilde{\Lambda}$. (a) An example of the PSM nuclei. (b) An example of the PSM nuclei following binarization. The red window illustrates the local average pixel intensity $\lambda$. (c) The probability distribution $P(\lambda)$ of the local average pixel intensity $\lambda$. The variance is a metric of the cell-scale texture $\Lambda$.

Here we give further details on the calculation of the normalized cell-scale texture $\tilde{\Lambda}$. The general process is outlined in Fig. S4. First, the nuclei data or simulation data is binarized. Then a window of side lengths $r$ similar to the size of a single cell is passed over the image, resulting in local average pixel intensities $\lambda = N_b/N_p$ where $N_b$ is the number of black pixels in the window and $N_p$ is the total number of pixels in the window. For each image there are $N$ sub windows and the cell-scale texture is then $\Lambda = \frac{1}{N}\sum_{i=1}^{N}(\lambda_i - \langle\lambda\rangle)^2$.

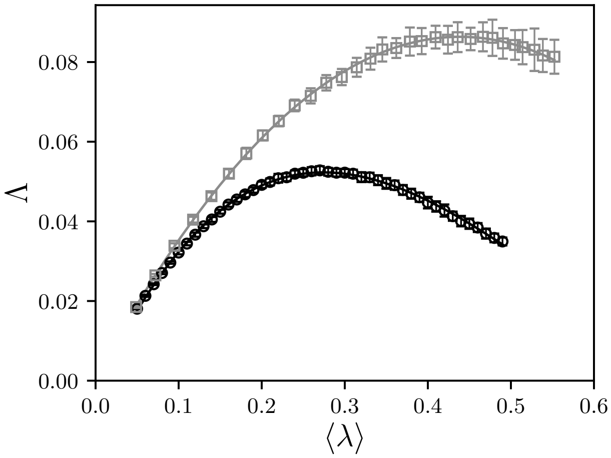


Figure S5: The particle-scale texture $\Lambda$ vs the average particle-scale fill fraction $\langle\lambda\rangle$ for RSA packings (black circles) and Poisson points (grey squares). Solid lines indicate 3rd degree polynomial fits to the two processes in black and grey respectively. The error bars indicate the standard deviation.

For a reference, we selected random sequential addition (RSA), where particles are placed randomly but without overlaps. In Fig. S5, we show how the value of $\Lambda$ varies for RSA packings as a function of $\langle\lambda\rangle$. We use the polynomial fit of this data to serve as a reference for the experimental data, whose $\langle\lambda\rangle$ varies slightly from image to image, and for the simulations where $\langle\lambda\rangle$ is fixed. Additionally, we compare the result for RSA to a Poisson point process. While Poisson points and RSA packings show the same scaling in texture vs length-scale, when looking at the particle length scale, the local textures are different. We selected RSA as the cells in both experiment and simulation are approximately non-overlapping.

To measure the structure of the tissue images, we measure the variance in the texture $\Lambda(r)$ using a box size $r$ approximately the size of the cells of $r \sim 12$ $\mu$m. Additionally, we normalize the variance to the expected texture variance for a random packing of non-overlapping disks at the same fill fraction by measuring $\Lambda(r)$ with a box size equal to the average particle size $r = 1.2$. While the average cell size is only approximate, we demonstrate in Fig. S6 that the normalized texture $\tilde{\Lambda}$ of the nuclei labeled PSM is robust to box size.

The particle scale texture can also be measured for the cytoplasmically labeled PSM. In Fig. S7 (a) and (b) we present an example image of the cytoplasmically labeled PSM before and after applying the binarization protocol used in the main text respectively. Calculating cell-scale texture in the same way and normalizing by RSA results in $\tilde{\Lambda} = 0.955 \pm 0.146$, a larger value than seen for the nuclei, but still as uniform as the uniform random distribution. (See Fig. S7 (c)). However, as the cytoplasmic label labels the entire cell, and the cellular architecture in the PSM is a interconnected network structure, RSA may not be the appropriate reference structure. Poisson placed points, where overlaps are allowed, have a similar level of connectivity as the cytoplasmically labeled data. (See Fig. S7 (d)). When normalizing by Poisson, we find $\tilde{\Lambda} = 0.432 \pm 0.079$ suggesting that for mass that is as interconnected as the PSM cytoplasmically labeled images are, the tissue still appears quite uniform.

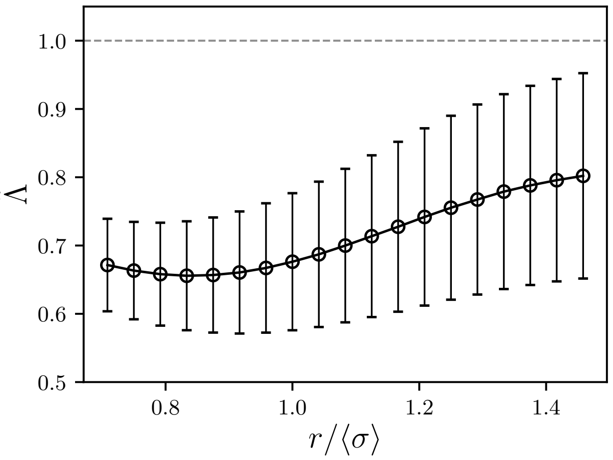


Figure S6: Average local texture of the nuclear labeled PSM vs the box size relative to the average particle size $r/\langle\sigma\rangle$ normalized to RSA disk packings at the same fill fraction. Error bars indicate the standard deviation.

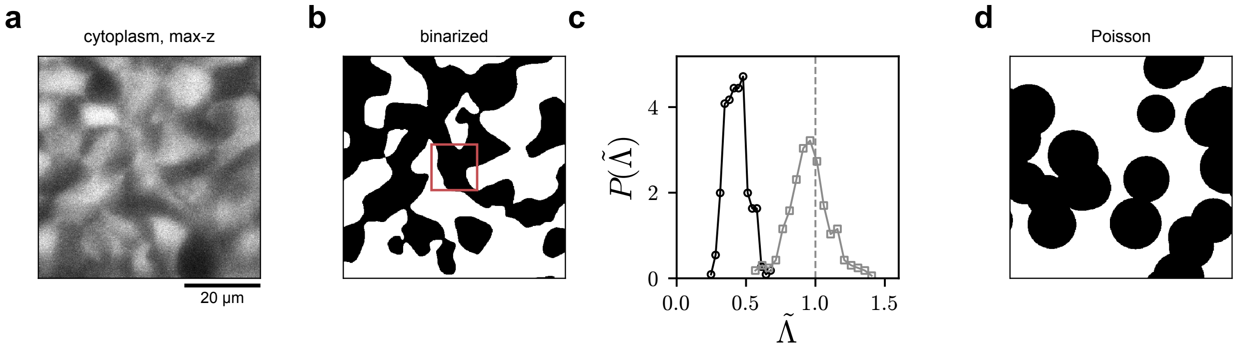


Figure S7: Texture analysis on cytoplasmically labeled PSM. (a) Example max-z projection of the cytoplasmically labeled PSM. (b) The same image following binarization. The red box indicates the window size used at the approximate cell size. (c) Probability distributions of the normalized cell-scale texture $\tilde{\Lambda}$ of the cytoplasmically labeled PSM using Poisson (black circles) vs RSA (grey squares) as the normalizing reference pattern. (d) An example of a Poisson process of random, uniformly placed overlapping disks at the same fill fraction.

Lastly, since nuclei are slightly ellipsoidal and therefore not perfectly isotropic, we consider the effect of particle shape anisotropy on the texture variance. In Fig. S8, we show example RSA packings of polydisperse ellipses with aspect ratio $\alpha$ and plot the texture variance $\Lambda$ vs the average particle scale texture $\langle\lambda\rangle$, using a box size equal to the diameter of a circle with the average ellipse area. Overlaps between ellipses are detected exactly using the Perram–Wertheim contact function [1]. While anisotropy reduces the texture variance by about 10% at the highest values of $\alpha \sim 2$, nuclei are typically fit by ellipses with $\alpha \sim 1.2$, which changes $\Lambda$ by less than 1%. Given this small effect, for simplicity, we use the isotropic RSA patterns for normalization.

Taken together, the cytoplasmic labeling result and the small effect of aspect ratio supports the conclusion that the PSM cells are more uniformly dispersed than randomly placed and clustered objects.

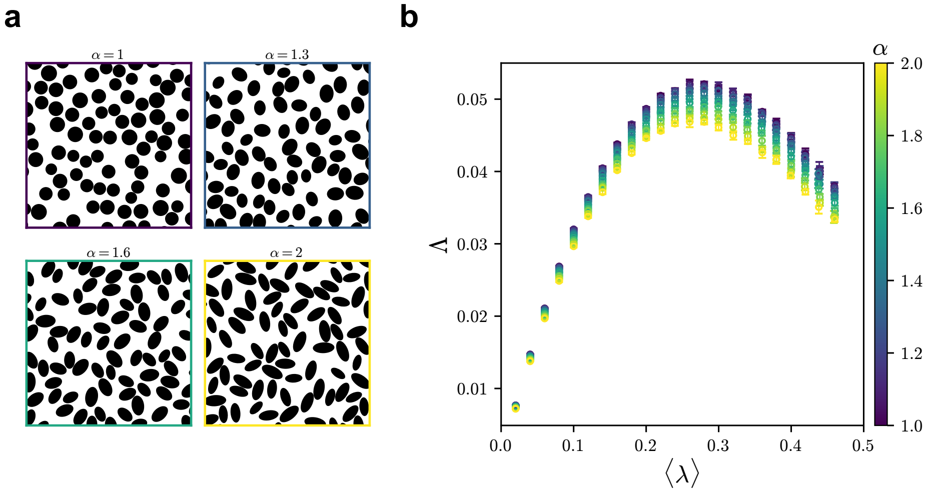


Figure S8: (a) Example RSA ellipse packings at fill fraction $\langle\lambda\rangle = 0.42$ with aspect ratio $\alpha = 1$, 1.3, 1.6 and 2, outlined in the color of their $\alpha$. (b) The particle scale texture $\Lambda$ vs the average fill fraction $\langle\lambda\rangle$ for RSA ellipse packings colored by aspect ratio $\alpha$ increasing from purple to yellow. Error bars indicate the standard deviation.

### 4 Scattering angle of non-interacting Active Brownian Particles

In the main text, we show that free crawling cells that do not collide have a scattering angle of $\chi \sim 34°$. Here we show that the origin of this scattering angle is what one would see for ideal non-interacting active Brownian particles (ABPs). First, we generate ideal non-interacting ABP trajectories across a range of rotational diffusion coefficients $D_r$. Then, we extract the scattering angle across a time window $w$ as done in the main text, plotted against the dimensionless product $wD_r$, resulting in the grey line in Fig. S9. In the main text, for computing scattering angles we used a window of ±30 minutes on either side of the collision, which corresponds to the X in Fig S9. Given longer windows, or faster rotational diffusion, the relative velocity vectors into and out of the window become uncorrelated and reach the expected angle between two random vectors of 90°. Additionally, we extract the apparent $D_r$ from the experimental non-colliding cell tracks and calculate the resulting $\chi$ resulting from different window times $w$ (black circles). We find the experimental data to lie on top of the numerical ABP data, indicating that the non-colliding cells scatter as one would expect if they were crawling like non-interacting ABPs. A plateau in $\chi$ occurs at small $w$ likely due to the time resolution in the data. Additionally, we were not able to extract a significant thermal diffusion on top of the rotation diffusion $D_r$ and therefore simulated the ABPs under the limit of $T = 0$.

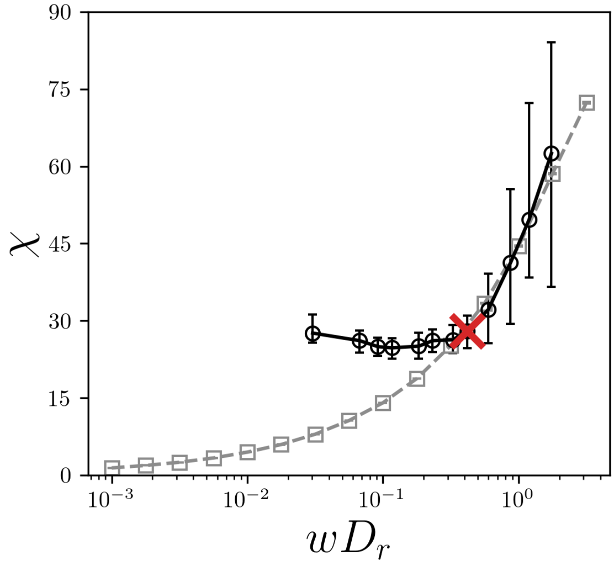


Figure S9: The scattering angle $\chi$ vs the observation window time $w$ scaled by the rotational diffusion constant $D_r$ for numerically simulated non-interacting active Brownian particles (grey squares) and extracted from non-interacting single cell tracks (black circles) from the explant spreading experiments. The red x indicates the time window used to calculate the scattering angle for the experimental data in the main text.

### 5 Void Analysis

How should a void be defined so that its growth reflects the tissue rather than the analysis? Two parameters set what counts as a void: the minimum void size and the morphological closing used to find voids.

First, a void is a region where cells are missing, so we set the minimum void size in units of cells rather than pixels. We retain only cleared regions larger than half the area of a 10 $\mu$m-diameter cell (39 $\mu m^2$) to avoid oversegmenting potential noise in the image and not true persistent voids. Second, finding the explant envelope and finding voids require opposite amounts of smoothing. The envelope needs a strong closing (a disk of radius 3 pixels) so that gaps between cells at the edge do not let the boundary leak inward, but the same closing fills in small voids. We therefore find voids in a separate mask closed with a disk of radius 1 pixel, the weakest closing that removes single-pixel noise, and clip it to the envelope.

In Fig. S10, we show four frames of one explant without (top row) and with (bottom row) the extracted voids, using the same frames, segmentation and intensity scale as Fig. 4 (g) of the main text. The labeled regions show strong visual agreement with the dark, cell-free regions of the unlabeled images, and the void area grows in the sequence described in the main text: no voids at $t = 0$, small voids between departing cells and the new cluster edge at 5 hours, the first interior void at 6 hours, and large central voids by 8 hours.

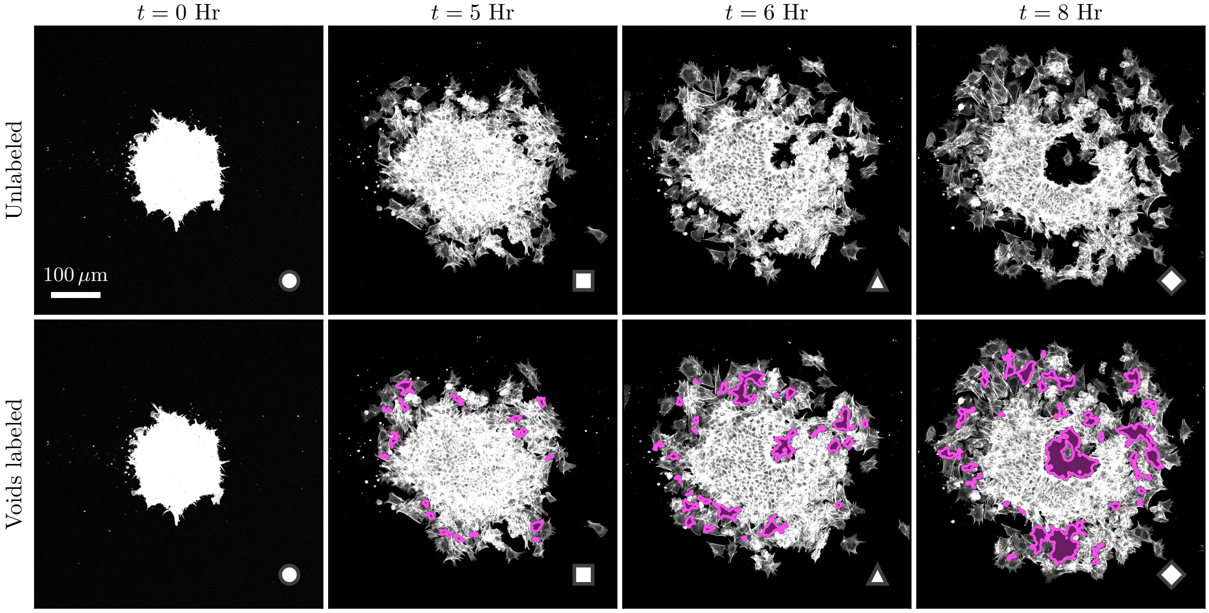


Figure S10: Void segmentation of the explant in Fig. 4 (g) of the main text. (Top row) Maximum-intensity projections at $t = 0$, 5, 6 and 8 hours, without annotation. (Bottom row) The same frames with the extracted voids shaded in pink. Markers match those in Fig. 4 (g, h). Each frame is divided by its mean intensity and displayed on the same gray scale in both rows, saturating at four times the mean, which saturates the dense cluster at $t = 0$. All panels share one crop box, centered on each frame's center of mass. Scale bar, 100 $\mu$m.

### 6 Further numerical CIL results

Here, we present further data on the tensioned fluid phase in both models. First, in Fig. S11, we present mean-squared displacements of both CIL-C and CIL-P models in the tensioned fluid phase for $k_{\text{off}} = 10^{-5}$ for both models and $v_0 = 5 \times 10^{-2}$ for the CIL-C and $k_r = 10^{-3}$ and $k_{\text{on}} = 10^{-3}$ for the CIL-P. The linear scaling shows that both models in the tensioned fluid phase can be diffusive. Next, in Fig. S12, we present visualizations of the three different phases both CIL models take with increasing activity: frozen, clustered and tensioned fluid. With little activity, either through crawling or ratcheting, and high unbinding, both models relax their initial network quickly until they become frozen. In the case of CIL-C, which only crawls if an adhesion is present and only makes new adhesions on contact, with a fast unbinding and slow crawling, all contacts are lost and the system freezes. In the case of CIL-P, the high on rate still adds Voronoi neighbors as contacts, shown as long range cell-cell contacts, however; the ratchet rate becomes slow enough relative to the fast off rate that very little energy is injected into the system and it also stops moving. As activity is increased relative to the adhesion life time, a clustered phase emerges. This occurs as the springs relax faster than the system generates tensions, causing cells to come together and cluster. Finally, with enough activity to overcome relaxation, a persistent, rearranging, tensioned, system spanning network is formed.

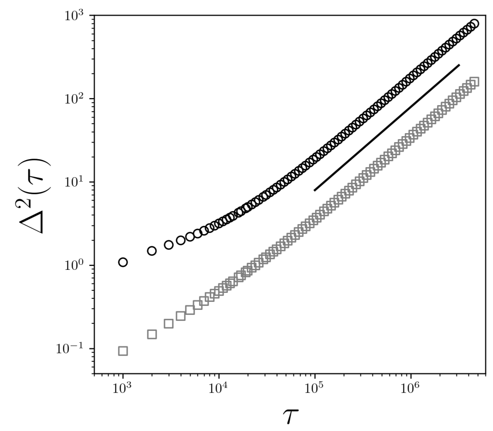


Figure S11: Mean-squared-displacement vs lag-time for the CIL-C model (black circles) and CIL-P model (grey squares) for $k_{\text{off}} = 10^{-5}$ for both models, $v_0 = 5 \times 10^{-2}$ for CIL-C and $k_r = 10^{-3}$ and $k_{\text{on}} = 10^{-3}$ for CIL-P. The black line indicates the diffusive slope of 1.

In the main text, we present the CIL-P results using a constant on rate $k_{\text{on}} = 10^{-3}$. Here, we present the same results but for $k_{\text{on}} = 10^{-4}$ and $k_{\text{on}} = 10^{-5}$ as well shown in Fig. S13. Overall, the results persist. However, as the on rate decreases, the amount of time available to the system to relax increases, leading to deviations from the theoretical predictions in the low activity region. For example, for $k_{\text{on}} = 10^{-5}$, below the plateau in tension and diffusion, the peak in diffusion increases and there is less tension than predicted. The tension prefactor is unchanged, $\mathcal{T} \approx 1.3\, k\langle c\rangle l_0$ on the plateau for all three on rates, but the tension crossover moves from $k_r/k_{\text{off}} \approx 10$ to 36 and 176 for $k_{\text{on}} = 10^{-3}$, $10^{-4}$ and $10^{-5}$, and the tension at fixed $k_r/k_{\text{off}}$ spreads with $k_{\text{off}}$.

Next, we test whether cell division and death disrupt the tensioned fluid. In the simulations above the number of cells is fixed, but dividing cells are visible in the PSM movies. We add cell division by inserting a daughter cell in contact with its parent at a random angle, and cell death

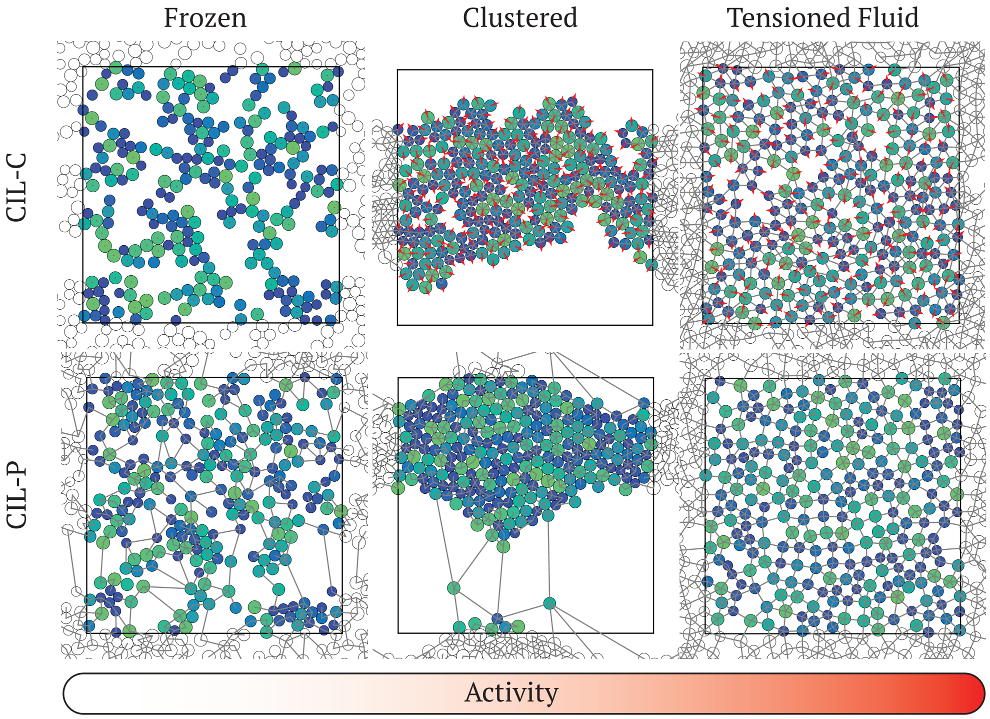


Figure S12: Visualizations of (top row) CIL-C (bottom row) CIL-P in the (left column) frozen, (center column) clustered and (right column) tensioned fluid phases. Cell color indicates the repulsive diameter increasing from blue to green. Red arrows in CIL-C indicate crawling direction. Grey lines indicate adhesions.

by removing a cell together with all of its adhesions. Every $10^3$ time units, each cell divides with probability $p_b(N_0/N)^4$ and dies with probability $p_d(N/N_0)^4$, where $N$ is the current number of cells, $N_0 = 256$ is the target number and $p_b = p_d = 10^{-4}$, so that the population fluctuates about homeostasis and each cell is replaced about once over a simulation. In Fig. S14, we compare the CIL-P model with and without cell division and death for $k_{\text{on}} = 10^{-4}$, $k_{\text{off}} = 10^{-5}$ and $k_r = 10^{-3}$. The number of cells stays within 4% of $N_0$, the mean-squared displacement remains diffusive with a diffusion coefficient within 10% of the value at fixed $N$, and the average tension per adhesion is unchanged, $\mathcal{T} \approx 0.84$ in both cases, although it fluctuates more. The texture rises slightly, from $\tilde{\Lambda} \approx 0.87$ to 0.93, while remaining more uniform than random. Fig. S15 shows snapshots of recently added and removed cells. We find the same behavior for the CIL-C model (data not shown). The tensioned fluid is a nonequilibrium steady state that recovers from the local perturbations of cell division and death in the same way that it resists the clustering instability, as new adhesions continually form toward the gaps that removed cells leave behind.

Could tension imposed from outside the tissue, for example by body-axis elongation via extracellular swelling, replace the tension that the cells generate locally? To test this alternative, we take a CIL-P network in the tensioned fluid phase (Fig. S16 (a)) and slowly stretch it along $x$ to twice its width, once with ratcheting activity and once without. Without ratcheting, adhesions still bind and unbind, but they carry tension only from the imposed extension.

In both cases the extension puts the network under tension. However, without ratcheting the cells gather into dense clumps separated by large gaps (Fig. S16 (b)), while with ratcheting they

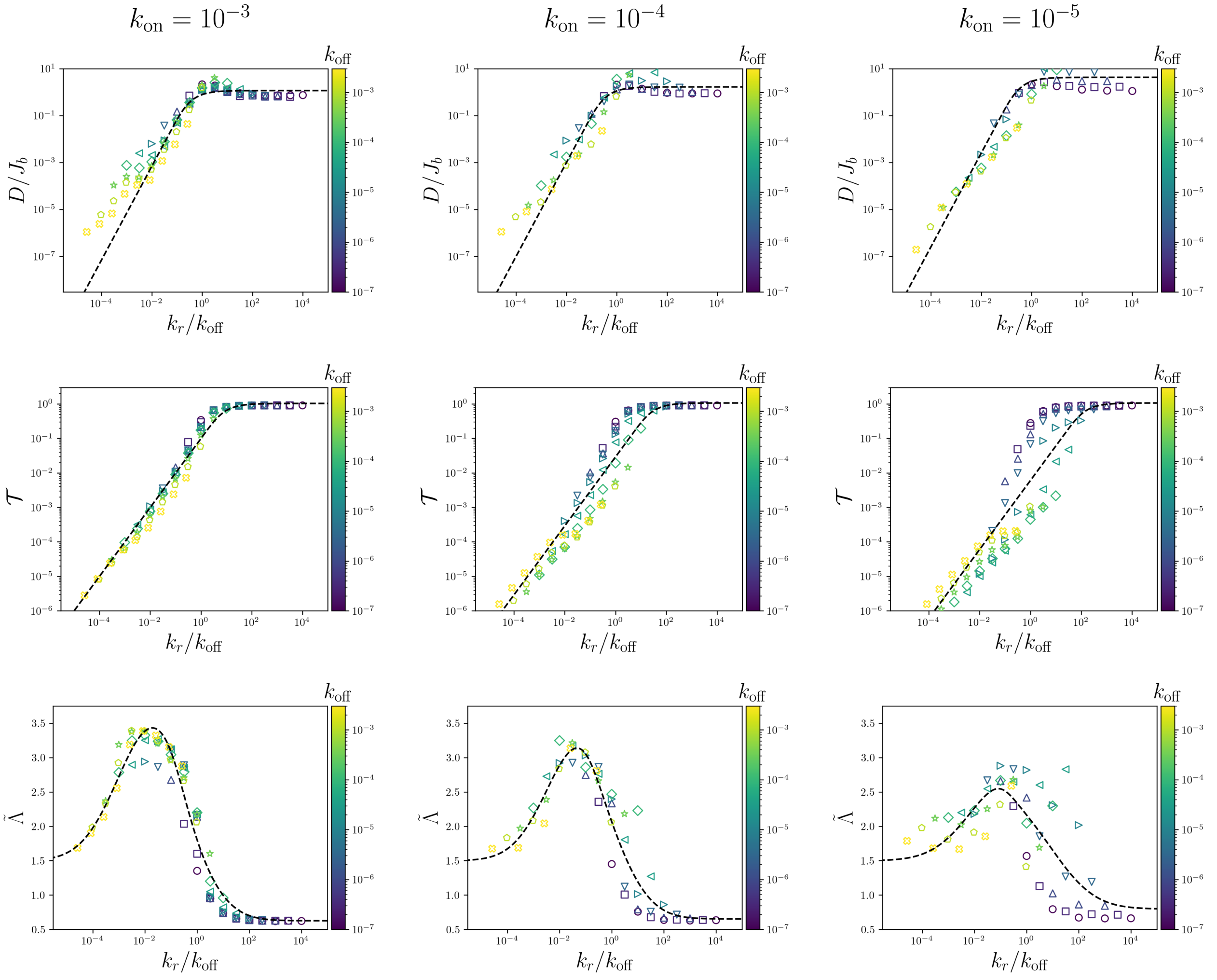


Figure S13: (Top row) The diffusion coefficient relative to the binding flux $D/J_b$, (middle row) the tension per adhesion $\mathcal{T}$ and (bottom row) the normalized cell-scale texture $\tilde{\Lambda}$ vs the ratchet rate relative to the off rate $k_r/k_{\rm off}$. Black dashed lines indicate $D/J_b \sim \langle c \rangle^2$, $\mathcal{T} \sim k\langle c \rangle l_0$ and Eq. 7 of the main text, each with one fitted prefactor per column; the tension crossover is also fitted per column. Color indicates increasing $k_{\rm off}$ from purple to yellow. On rates $k_{\rm on}$ are indicated by the column labels.

remain uniformly dispersed across the doubled box (Fig. S16 (c)). The clumping is the clustering instability described in the main text: as the stretched network rearranges, gaps open between cells, and without local contraction nothing pulls those gaps closed, so they grow. Thus a global extension alone does not generate the uniform texture of the PSM, which requires tension generated locally by the cells. Although this is a simulation result, the same logic applies to the embryo. If tension were applied only at the tissue boundary, the neighbor exchanges we observe would open holes that could not close, whereas holes close in the PSM movies. This test does not exclude a role for the surrounding tissues, however. To generate tension the cells must pull against something, and adhesion to neighboring tissues or to the extracellular matrix likely provides that anchoring.

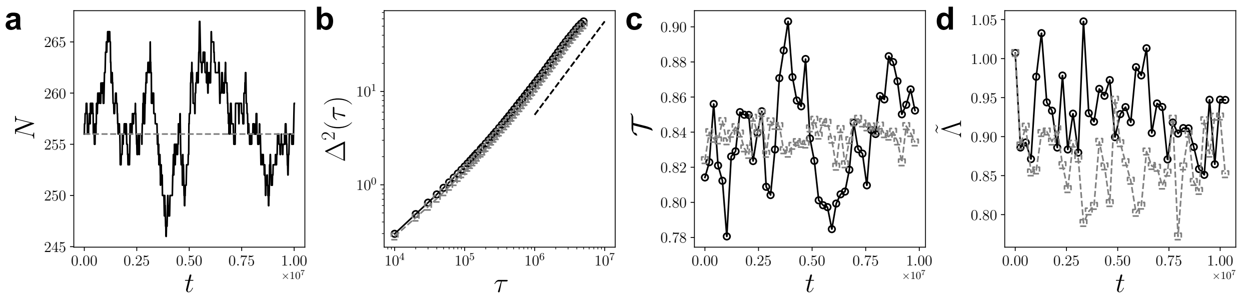


Figure S14: Comparison of the CIL-P model with (black solid lines, circles) and without (grey dashed lines, squares) cell division and death for $k_{\text{on}} = 10^{-4}$, $k_{\text{off}} = 10^{-5}$ and $k_r = 10^{-3}$. (a) The number of cells $N$ vs time. (b) The mean-squared displacement $\Delta^2(\tau)$ vs lag-time $\tau$. The dashed line indicates a diffusive slope of 1. (c) The tension per adhesion $\mathcal{T}$ vs time. (d) The normalized cell-scale texture $\tilde{\Lambda}$ vs time.

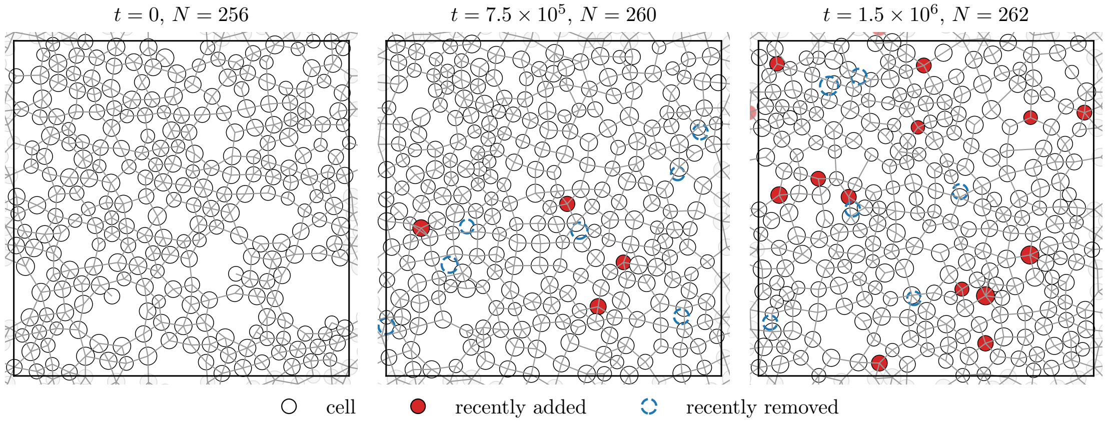


Figure S15: Snapshots of the CIL-P model with cell division and death at the times and cell numbers indicated. Recently added cells are shown in red and recently removed cells as blue dashed circles. Grey lines indicate adhesions.

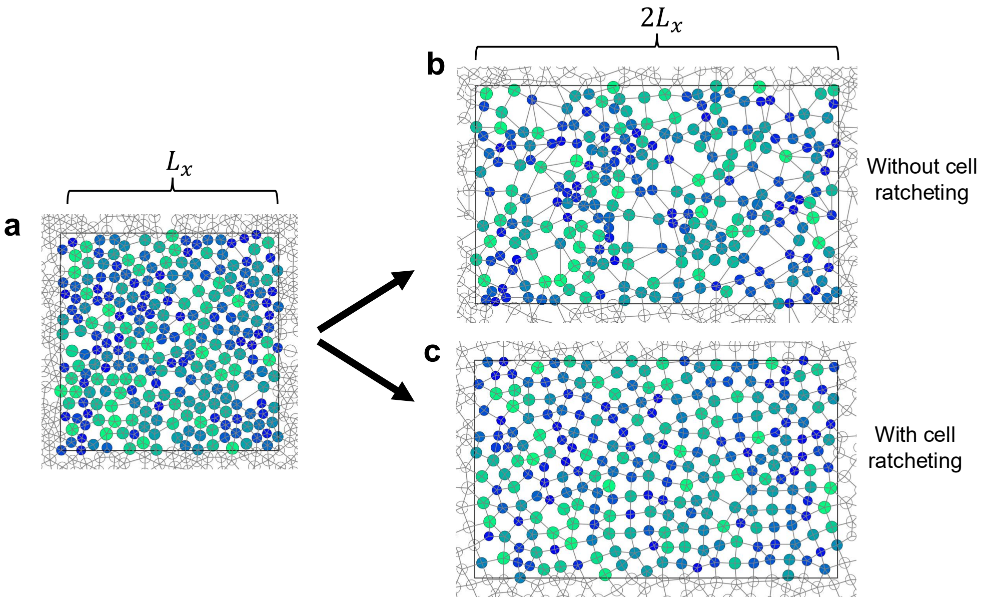


Figure S16: Global extension of a CIL-P network with and without ratcheting. (a) Initial configuration in the tensioned fluid phase, with box width $L_x$. (b, c) The same configuration after the box is stretched along $x$ to $2L_x$, (b) without ratcheting, $k_r = 0$, and (c) with ratcheting, $k_r = 10^{-4}$. Cell color indicates the repulsive diameter, increasing from blue to green, and grey lines indicate adhesions. The box is stretched over about six adhesion lifetimes $1/k_{\text{off}}$, and $\phi$ decreases from 0.6 to 0.3; $k_{\text{on}} = 10^{-3}$, $k_{\text{off}} = 10^{-5}$ and $N = 256$.

### 7 Further analytic CIL results

Here, we provide further rationale for the analytic predictions for both CIL-C and CIL-P models. First, we derive the tension scaling shared by both models. We start from the virial stress tensor,

$$\sigma_{\alpha\beta} = \sum_{\langle i,j \rangle} F_{ij}^{\alpha} r_{ij}^{\beta} \tag{S1}$$

where $\alpha, \beta \in \{x, y\}$, $F_{ij}^{\alpha}$ is the $\alpha$-component of the force between cell $i$ and $j$ and $r_{ij}^{\beta}$ is the $\beta$-component of the separation vector between cell $i$ and $j$. As we define the tension as the negative virial pressure per adhesion (see Methods),

$$\mathcal{T} = \frac{2L^2}{N_b}\left(-\frac{\sigma_{xx} + \sigma_{yy}}{2L^2}\right) = -\frac{1}{N_b}\sum_{\langle i,j \rangle} \vec{F}_{ij} \cdot \vec{r}_{ij}, \tag{S2}$$

an adhesion carrying a tensile force $F_b$ over a separation $r_b$ contributes $\mathcal{T} \approx F_b r_b$. In CIL-P, the rest length contraction is converted into spring extension, so $F_b = k\langle c \rangle$, and adhesions span neighboring cells, $r_b \sim l_0$. In CIL-C, a crawling cell held by its adhesions loads them with $F_b \sim \gamma v_0$, and adhesions form on contact, $r_b \sim \langle \sigma_{ij} \rangle$. The number of adhesions does not enter $\mathcal{T}$. Instead, two-state kinetics sets it, $N_b = N z_{max} f_b/2$, so that the total load carried by the network scales as $N_b \mathcal{T} \propto f_b F_b r_b$. In Fig. S17, we test this for CIL-P and find that the measured number of adhesions per cell follows $z = z_{max} f_b$ to within 0.1 decades for $k_{\text{on}} = 10^{-3}$, where $f_b$ spans 0.25–1. At lower on rates and small $f_b$, adhesions that form on contact, which the kinetic theory ignores, raise $z$ above $z_{max} f_b$.

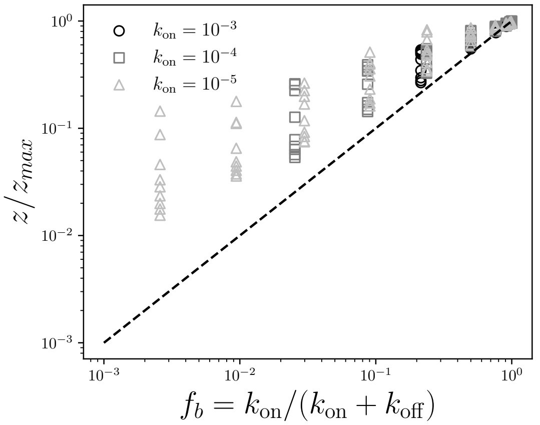


Figure S17: The measured number of adhesions per cell relative to its maximum, $z/z_{max}$ with $z_{max} = 4.8$, vs the two-state fraction bound $f_b = k_{\text{on}}/(k_{\text{on}} + k_{\text{off}})$ for CIL-P with $k_{\text{on}} = 10^{-3}$ (black circles), $10^{-4}$ (grey squares) and $10^{-5}$ (light grey triangles). The dashed line indicates $z/z_{max} = f_b$.

Moving to the CIL-P model, we can use the kinetic nature of the model to estimate the time average rest length contraction distance $\langle c \rangle$ analytically. First, we write the time average contraction in terms of the model parameters,

$$\langle c \rangle = \int_0^{\infty} c(t) P(t) dt \tag{S3}$$

where $c(t) = k_r \sigma_{ij} t$ is the linear contraction with rate $k_r$ until reaching the maximum contraction distance at $t_{max}$ and then is a constant value of $c(t_{max})$, and $P(t) = k_{\text{off}} e^{-k_{\text{off}} t}$ is the exponential unbinding probability. Then, we split the integral into two terms,

$$\int_0^{\infty} c(t)P(t)dt = k_r \sigma_{ij} k_{\text{off}} \int_0^{t_{max}} t e^{-k_{\text{off}} t} dt + c(t_{max}) k_{\text{off}} \int_{t_{max}}^{\infty} e^{-k_{\text{off}} t} dt \tag{S4}$$

representing the two stages of first contraction, followed by no contraction, all with a probability $P(t)$ to unbind. The initial spring length assuming uniformly distributed cells is $l_0 = L/\sqrt{N}$. With a minimum contraction distance set by the contact distance $\sigma_{ij}$, the maximum amount of contraction is $c(t_{max}) = l_0 - \sigma_{ij}$. The linear contraction rate $k_r$ then results in $t_{max} = (l_0 - \sigma_{ij})/(k_r\sigma_{ij})$. Both integrals are explicitly solvable and through simplification results in the final expression,

$$\langle c\rangle = \sigma_{ij}\frac{k_r}{k_{\text{off}}}(1 - e^{-k_{\text{off}}t_{max}}). \tag{S5}$$

Using $F_b = k\langle c\rangle$ and $r_b \sim l_0$ in the virial estimate above gives $\mathcal{T} \sim k\langle c\rangle l_0$. On the plateau we find $\mathcal{T} \approx 1.25\, k\langle c\rangle l_0$. However, the tension saturates at $k_r/k_{\text{off}} \approx 10$ for $k_{\text{on}} = 10^{-3}$, later than $\langle c\rangle$, which saturates at $(l_0 - \sigma_{ij})/\sigma_{ij} \approx 0.4$. We fit this crossover rather than derive it (dashed line in Fig. 5(b) of the main text), and it moves to larger $k_r/k_{\text{off}}$ as $k_{\text{on}}$ decreases (Fig. S13). The crossover coincides with the loss of clustering in the texture (Fig. 5(c) of the main text), which suggests that below it part of the contraction is balanced by compressed contacts between cells within clusters rather than stored in the adhesion network. Testing this requires separating the adhesion and contact contributions to the virial, which we leave for future work. We also tested $k_r/k_{\text{on}}$, $k_r/(k_{\text{on}} + k_{\text{off}})$, $k_r/J_b$ and $k_r/\sqrt{k_{\text{on}}k_{\text{off}}}$ as collapse variables across all three on rates, and $k_r/k_{\text{off}}$ remains the best (rms 0.70 decades against 1.2–1.8 decades). Similarly, if each binding event displaces a cell by $\sim \langle c\rangle$ in a random direction, the cells undergo a random walk with step size $\langle c\rangle$ at the binding flux $J_b$, and $D \sim \langle c\rangle^2 J_b$. Numerical factors of order one, such as the number of adhesions per cell and correlations between successive steps, are absorbed into the $\sim$ relations. Both relations are confirmed by the collapse presented in the main text.

Finally, we turn to the kinetic theory for the CIL-C model. The theory follows the same steps as for CIL-P with one change: $k_{\text{on}}$ is not an input parameter. A CIL-C cell only crawls while it holds an adhesion, and new adhesions only form when crawling closes a gap between cells. We therefore assume that crawling-driven contact formation is a first-order process with a rate proportional to the crawling force, $k_{\text{on}} = \beta\gamma v_0$. The two-state kinetics of the main text then carry over directly,

$$f_b = \frac{X}{1+X}, \qquad J_b = k_{\text{off}} f_b, \qquad X = \frac{k_{\text{on}}}{k_{\text{off}}} = \frac{\beta\gamma v_0}{k_{\text{off}}}, \tag{S6}$$

where we measure $f_b = z/z_{max}$ from the average number of adhesions per cell $z$. When the crawling extension of an adhesion $\delta = \gamma v_0/k$ is small, the network self-organizes to isostaticity, $z_{max} \approx z_c = 4$ in two dimensions, and $z_{max}$ grows as $z_c + a\sqrt{\delta}$ once the crawling force stretches adhesions (Fig. S18(c)). The collapse of $z$ gives $\beta \approx 3.5 \times 10^{-3}$, and the effective $\beta$ is flat across four decades of $\gamma v_0$ (Fig. S18(a)).

The frozen phase follows directly from these kinetics. As $z = 0$ is an absorbing state in CIL-C, the network falls apart once unbinding outpaces crawling-driven binding, $X \lesssim 1$, or $\gamma v_0 \lesssim k_{\text{off}}/\beta$, shown as the dashed line in Fig. S18(b).

For diffusion, the distance a released cell crawls before a new adhesion catches it, $\ell_D$, plays the role of the spring contraction in CIL-P. A released cell moves as $c(t) = vt$ until it has crossed $\ell_D$, and crossing $\ell_D$ is forming the new contact, so $v = k_{\text{on}}\ell_D$. The same integral as in Eq. S5 then gives

$$\langle c\rangle = \ell_D X\left(1 - e^{-1/X}\right), \tag{S7}$$

and, as for CIL-P, $D/J_b \sim \langle c\rangle^2$. The step $\ell_D$ depends on the structure of the network. In the clustered phase ($\delta \ll g$, where $g = l_0 - \langle\sigma_{ij}\rangle \approx 0.49$ is the average gap between uniformly spaced cells), cells cross the voids between clusters and $\ell_D = \ell_{cl}$. In the tensioned fluid ($\delta \gtrsim g$), the crawling extension spans the gap, the network is uniformly dilated, and each adhesion turnover moves a cell by one neighbor spacing, $\ell_D = l_0$. We interpolate between the two as $\ell_D^2 = l_0^2 + (\ell_{cl}^2 - l_0^2)e^{-\delta/g}$ and find $\ell_{cl} \approx 2.4\, l_0$ (Fig. S18(d)).

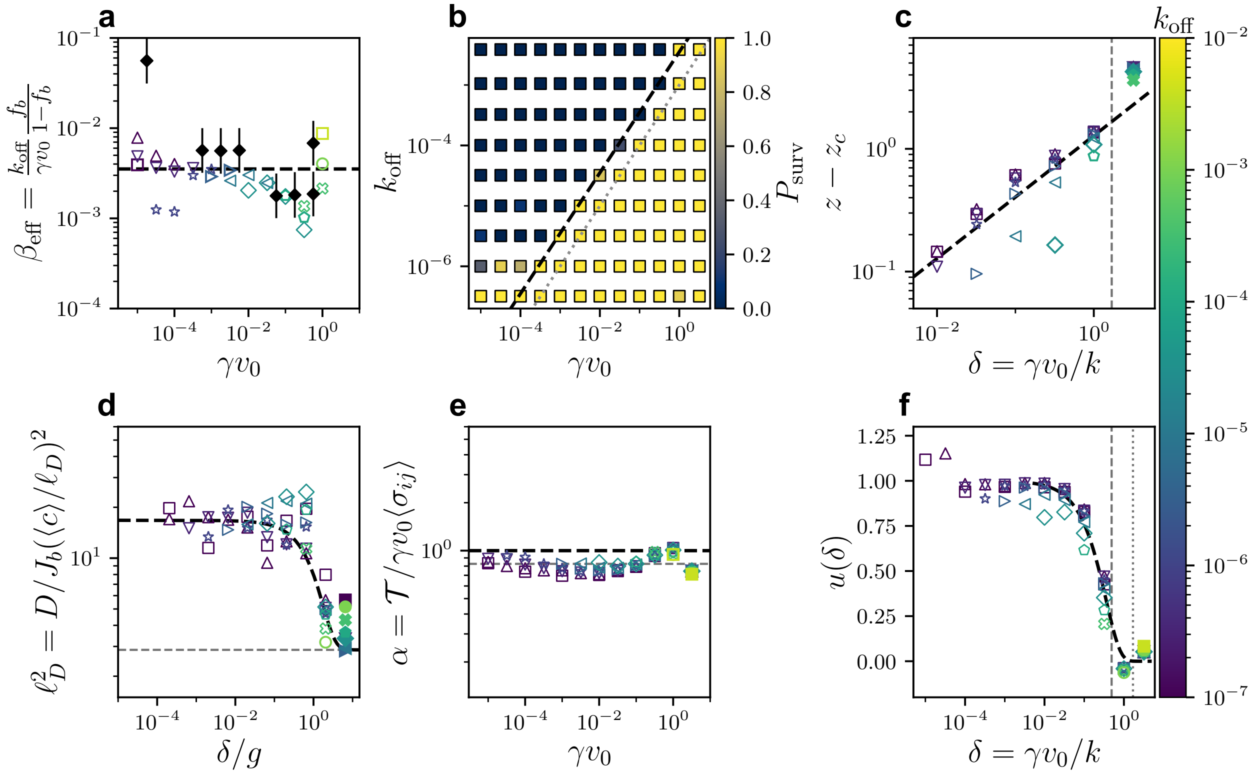


Figure S18: Inputs to the CIL-C theory. (a) The effective binding constant $\beta_{eff} = (k_{\rm off}/\gamma v_0)f_b/(1-f_b)$ vs $\gamma v_0$. The dashed line indicates the fit $\beta$ and black diamonds indicate the independent estimate from the frozen boundary. (b) The fraction of replicates that keep a spanning network $P_{surv}$ over $\gamma v_0$ and $k_{\rm off}$. The dashed line indicates $k_{\rm on} = k_{\rm off}$ and the grey dotted line indicates $\gamma v_0 = k_{\rm off}$. (c) The excess coordination $z - z_c$ in the saturated regime vs $\delta = \gamma v_0/k$. The dashed line indicates slope 1/2. (d) The step length $\ell_D^2$ vs $\delta/g$. The dashed line indicates the interpolation and the grey dashed line indicates $l_0^2$. (e) The tension per adhesion relative to the crawling force times contact distance, $\mathcal{T}/\gamma v_0\langle\sigma_{ij}\rangle$, vs $\gamma v_0$. The black dashed line indicates the force-balance value of 1 and the grey dashed line indicates the mean of the data, 0.83. (f) The texture switch $u(\delta)$ vs $\delta$. The dashed line indicates the random disk gap probability, and the grey dashed and dotted lines indicate $\delta = g$ and $\delta = l_0$.

For tension, a crawling cell held by its adhesions loads them with a force of order $\gamma v_0$, since the crawling direction $\hat{e}_i$ is normalized. Adhesions in CIL-C form on contact, so their separation is $\sim \langle\sigma_{ij}\rangle$, and the virial argument above gives $\mathcal{T} \sim \gamma v_0\langle\sigma_{ij}\rangle$, the CIL-C analog of $\mathcal{T} \sim k\langle c\rangle l_0$ for CIL-P. The measured tension per adhesion is $0.83 \pm 0.10$ of $\gamma v_0\langle\sigma_{ij}\rangle$ across four decades of $\gamma v_0$ (Fig. S18(e)).

For the texture, clustering grows with the bound fraction, and clusters persist only where crawling cannot span the gap to the nearest neighbor. For randomly placed disks, the probability that the nearest gap exceeds $\delta$ is $u(\delta) = \exp\left[-\pi n(2\langle\sigma_{ij}\rangle\delta + \delta^2)\right]$ (Fig. S18(f)). As an ansatz, we propose

$$\tilde{\Lambda} = \tilde{\Lambda}_t + \left[\tilde{\Lambda}_0 + (\tilde{\Lambda}_c - \tilde{\Lambda}_0)f_b - \tilde{\Lambda}_t\right] u(\delta), \quad \text{(S8)}$$

with $\tilde{\Lambda}_0 = 1$ for a random texture without adhesions, and we find the best fit $\tilde{\Lambda}_c \approx 3.9$ and

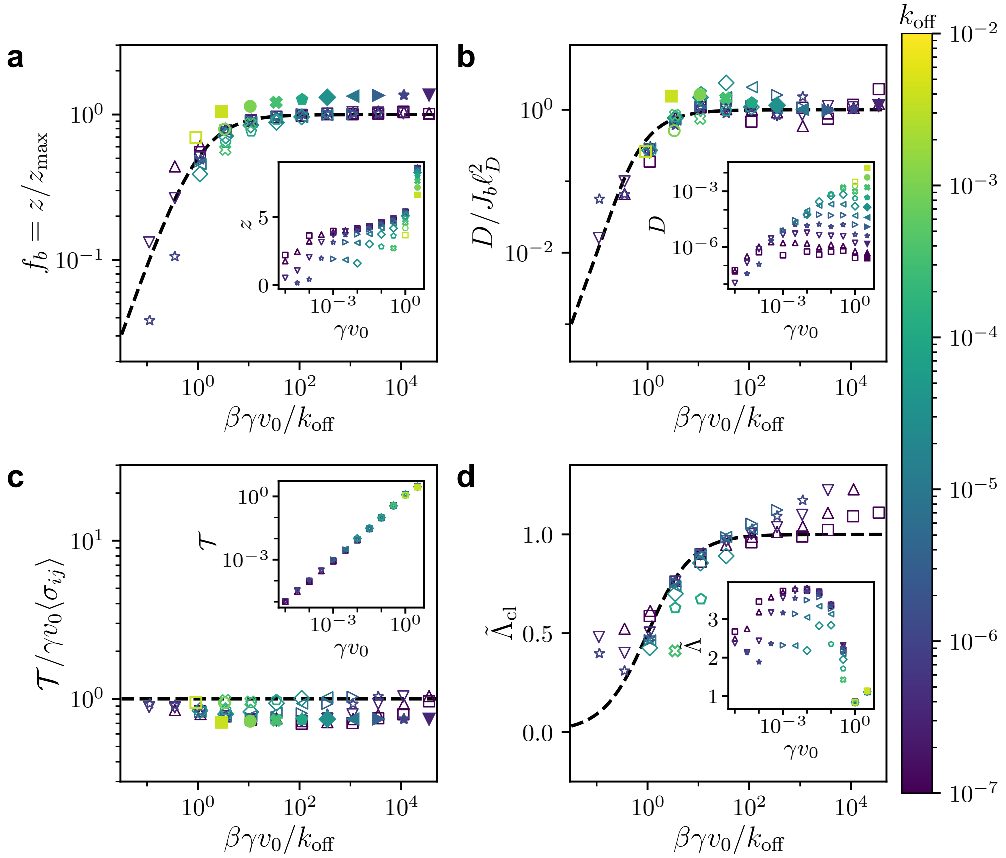


Figure S19: CIL-C collapse onto two-state kinetics with a crawling-driven on rate $k_{\rm on} = \beta\gamma v_0$. (a) The fraction bound $f_b = z/z_{max}$, (b) the diffusion coefficient $D$ relative to the binding flux $J_b$ and step length $\ell_D^2$, (c) the tension per adhesion $\mathcal{T}$ relative to $\gamma v_0\langle\sigma_{ij}\rangle$ and (d) the clustered branch of the normalized cell-scale texture $\tilde{\Lambda}_{cl}$ vs $k_{\rm on}/k_{\rm off}$. Dashed lines indicate $X/(1+X)$, $[X(1-e^{-1/X})]^2$, 1 and $X/(1+X)$ respectively. Open markers have $\delta < l_0$ and filled markers have $\delta \geq l_0$. Increasing unbinding rate $k_{\rm off}$ is colored from purple to yellow. Insets present the uncollapsed data.

$\tilde{\Lambda}_t \approx 0.96$.

In Fig. S19, we plot $f_b$, $D/J_b\ell_D^2$, $\mathcal{T}/\gamma v_0\langle\sigma_{ij}\rangle$ and the clustered branch of the texture vs $k_{\rm on}/k_{\rm off}$, and find that all four collapse onto the kinetic predictions with four fitted parameters ($\beta$, $\ell_{cl}$, $\tilde{\Lambda}_c$ and $\tilde{\Lambda}_t$). The theory breaks down in three places. First, next to the absorbing boundary ($z \lesssim 1.5$), mean-field kinetics fail, as expected near an absorbing-state transition. Second, for $\delta \approx 0.03$–$0.3$, $\ell_D^2$ still depends on $k_{\rm off}$ at fixed $\delta$ by $\pm 0.2$–$0.3$ decades. Third, for $\gamma v_0 = 3.2$, the crawling extension exceeds the neighbor spacing, cells crawl past their nearest neighbors, and $z$, $D$ and $\tilde{\Lambda}$ run above the predictions (filled markers), the regime noted in the main text Discussion. The texture of the slowest unbinding rates ($k_{\rm off} \lesssim 10^{-6}$) also overshoots the clustered branch by up to $\sim 20\%$ at $k_{\rm on}/k_{\rm off} \gtrsim 10^3$. Additionally, $\beta$ and $\ell_{cl}$ are fit rather than derived. Both are much larger than a single gap (the search length $1/\gamma\beta \approx 0.5L$), which suggests collective floppy-mode motion in the clustered network; if so, both should depend on $N$, while $\ell_D = l_0$ in the tensioned fluid should not.

### 8 Supplemental video captions

Video S1. Example of the nuclear labeled embryo, recorded over 2 hours. Lines indicate extracted continuous cell tracks, color from blue to red with increasing time.

Video S2. Example of the laser ablation study along with the extracted PIV velocity field. Cells are cytoplasmically labeled.

Video S3. Visualizations of all three activity models, SPP, CIL-C and CIL-P, in the top, middle and bottom rows respectively, for increasing activity from left to right. SPP undergoes a clustering transition. CIL-C and CIL-P undergo a re-entrant clustering transition before reaching the tensioned fluid phase. Color indicates increase cell size from blue to green. Grey lines indicate adhesions. Red arrows in SPP and CIL-C indicate the direction of activity.
